# Omega 3 fish oil suppress radiation induced hepato and renal toxicity in mice through modulation in Wnt canonical pathway combined with NHEJ and Intrinsic Apoptotic pathway

**DOI:** 10.1101/2023.02.05.527226

**Authors:** Shashank Kumar, Suttur S Malini

## Abstract

Radiation is associated with inflammation and oxidative stress, the latter of which contributes to activation of DNA damage and apoptosis. Omega 3 polyunsaturated fatty acids (PUFAs) have been reported to limit oxidative stress and DNA damage. The aim of this study was to evaluate the effect of the Omega 3 PUFA on antioxidant defence in male physiology on mice model. Liver and kidney tissues were obtained from whole body irradiated mice divided under 9 groups (Weight-10mg, 6-8 months old, n=5) and age-matched male controls (6-8 months old, n=5). 6 groups have been orally intubated with (50, 100 and 150) mg/kg BW with Omega 3 fish oil 1hr prior to the radiation exposure. Liver and kidney were surgically obtained after 24 hours and 30 days of radiation exposure. Omega 3 fish oil supplementation increased the level of mRNA expression of Lef1, Axin2, Survivin, Ku70, SOD1, SOD2, Cat, iNOS and decresed the level of Bax and Bcl2 in irradiated with omega 3 fish oil supplementation compare to irradiated alone. Omega 3 fish oil increased SOD scavenging, Catalase, Nitric oxide scavenging activity, Total antioxidant capacity and decrease the lipid peroxidation. The improvements in mRNA level of candidate genes of Wnt canonical pathway, NHEJ pathway, oxidative stress status serve as a stimulus for further investigation of Omega 3 fish oil as supplementation for patients undergo radiation therapy.

## Introduction

Docosahexaenoic acid (22:6n-3; DHA) and eicosapentaenoic acid (20:5n-3, EPA), two long-chain omega-3 polyunsaturated fatty acids (n-3 PUFAs), which usually present in fish oil have been proposed to have cardioprotective (Lee et al., 2008), anti-inflammatory (Layé et al., 2018), immunoregulatory (Alshatwi & Subash-Babu, 2018), antioxidant (Giordano & Visioli, 2014), and anti-tumor effects (Gevariya et al., 2019). These positive effects on human health have been linked to: (a) competition with arachidonic acid (AA) for enzymes involved in the biosynthesis of pro-inflammatory mediator molecules (Kwon, 2020); (b) suppression of pro-inflammatory nuclear factor kappa B (NF-kB) through modulation of toll-like receptor 4 (TLR4) signalling and activation of peroxisome proliferator-activated receptor gamma (PPAR gamma) (Calder, 2015); and (c) activation of the (FFA4, formerly GPR120) metabolism to pro-resolution lipid mediators (positions 11 and d) (e.g. resolvins, protectins, maresins)(Ferreira et al., 2022). Omega 3 PUFAs have recently been shown to have anti-inflammatory and pro-resolution effects in concert.to suppress pro-oxidant activity by upregulating genes encoding cytoprotective antioxidant proteins such as heme oxygenase 1 (HO-1) and glutathione peroxidase (GPx) (Ishikado et al., 2013; Kusunoki et al., 2013; Meital et al., 2019; Zhang et al., 2014). HO-1 is a single, transmembrane 32-kDa protein that plays a central role in stress adaptation(Origassa & Câmara, 2013). HO-1 provides cells and tissues with an inducible antioxidant defence mechanism that can be ubiquitously activated in response to elevated levels of heme, its natural substrate, and a multiplicity of endogenous factors such as heavy metals, cytokines, hormones, growth factors, nitric oxide and significant positive association between long chain polyunsaturated fatty acids (PUFAs) of the omega-3 series, eicosapentaeinoic acid (EPA)(Meital et al., 2019; Origassa & Câmara, 2013). Omega-3 FAs are considered as potential important antioxidants (Hodge et al., 2006). Recent studies have shown that Omega 3-PUFAs, DHA and EPA decreased the cell growth, and DHA exhibited more significant effect shown 80% growth inhibition at 50 μM concentration in breast cancer cell lines (Yun et al., 2016). Omega 6-PUFAs/Omega 3-PUFAs ratio in diets alter the risk of malignancy, though their exact patho-chemical 41 of 48 interactions with the tumor are still obscure. It has been reported that Omega 6-PUFAs stimulate carcinogenesis, tumor growth and metastasis, whereas Omega 3-PUFAs have inhibitory effects of them (Yun et al., 2016). DHA significantly had suppressed invasiveness by suppressing MMP-2 and MMP-9 secretion through inhibition of NF-κB and Cox-2 signaling pathway. The predominant mechanisms of DHA have been thought to be a reduction in pro-inflammatory eicosanoids and an increase in inflammation-resolving derivatives (Turk and Chapkin, 2013., Calder, 2004) Recent findings have suggested that Omega 3 polyunsaturated fatty acids (PUFAs) might be of use as adjuvant for cancer therapy. In addition to inhibiting the progression of several prevalent cancers (Pont et. al., 2006, Berquin et. al., 2008), Omega 3 PUFAs have been found to enhance the sensitivity of tumour cells to several anticancer drugs, such as mitomycin C, cyclophosphamide, and doxorubicin (Pardini 2001). On the other hand, Ionizing radiation is an important modality used in the treatment of malignancy and is one example of an agent that induces oxidative genotoxic stress. Ionizing radiation causes severe cellular damage and stress both directly, by energetic disruption of DNA integrity, and indirectly, as a result of the formation of intracellular free radicals (Kumar et al., 2022). A small number of studies have shown an effect of radiation on miRNA expression patterns both in vitro(Simone et al., 2009) and in vivo (Ilnytskyy et al., 2008; Josson et al., 2008; Koturbash et al., 2008). Interactions between the cellular ROS-producing systems are joined by ROS produced by water radiolysis. One of the earliest comprehensive descriptions of post-irradiation oxidative stress was given by Leach et al., 2001.

While it has been suggested that omega 3 PUFAs can reduce the negative effects of inflammation, oxidative stress, and disturbed antioxidant status in patient populations with radiation-related conditions, more research is needed to fully understand the therapeutic potential of these bioactive nutrients. The purpose of this study was to assess how Omega 3 fish oil affected antioxidant defence mechanisms as well as modulatory properties in the Wnt canonical pathway combined with NHEJ pathway and intrinsic apoptotic pathway.

## Materials and Method

### Group Distribution

50 adult Swiss albino mice (male) having an average weight of 25-30gm (Aged 6 to 8 months old) were taken for the study, and 10 groups were made in the arrangement of 5 mice per group for first experiment. Animals were divided into seven groups according to the Table 1

**Table 1.**

| Group | Autopsy at 24hrs | Autopsy at 30 days |
| --- | --- | --- |
| Control |  | 5 |
| Vehicle |  | 5 |
| Vehicle + Radiation | 5 | 5 |
| O3 fish oil + Radiation<br>(50mg/kg BW) | 5 | 5 |
| O3 fish oil + Radiation<br>(50mg/kg BW) | 5 | 5 |
| O3 fish oil + Radiation<br>(50mg/kg BW) | 5 | 5 |
|  | 20 | 30 |

### Irradiation

For irradiation, Siemens Linear Accelerator (HCG Bharat Cancer Hospital and institute of Oncology) was used. Each mouse was kept in cage and fixed at the distance of 12cm from the accelerator. Doses were pre calibrated according to the Institution guidelines. Swiss albino mice were kept in a cage made up of polypropylene 4×4×4 cm3 cage. These animals were irradiated at 6 Mev to radiation dosages according to Table. 1.

### Preparation of Sample

Mice were sacrificed with the fulfilment of ethical parameters. liver, spleen, and kidney (Fig. A) were removed and kept in PBS with pH-7.4. 10mg of tissue from each group were homogenized with 1ml of PBS and kept centrifuged at 3000 rpm for 10 minutes. The supernatant was used to evaluate biochemical parameters.

## Evaluation of Oxidative Stress Parameters

### Superoxide Scavenging Activity

Superoxide dismutase activity has been estimated by the modified method of (Kazari Das et al, 2000). The 100 μl supernatant of homogenized tissue in PBS was taken as the sample. The reaction mixture, phosphate buffer (0.1 M pH-7.4), α-methionine (20 mM), Hydroxyl Amine Hydrochloride (10 mM), EDTA (50 μM), Triton-X (1%), Riboflavin (100 μM) were added to the sample. Without sample serves as a control and without riboflavin serves as blank and the spectrophotometer observed the OD.

*Scavenging effect %= [Absorbance of control/ Absorbance of sample - 1] ×100*

## Catalase activity

Catalase (Cat) activity was measured by adding 50 μl of the supernatant to a 3 ml reaction mixture containing 8.8 mM H_2_O_2_ (1.5%) and 0.1 M sodium phosphate buffer (pH-7.4), and the absorbance at 240 nm was monitored for 3 min. The decrease in H_2_O_2_ concentration was expressed as mmol of H_2_O_2_ decomposed/min/mg of protein (Aebi, 1984)

## Lipid peroxidation

Lipid peroxidation (LPO) in the homogenate was measured by the thiobarbituric acid reactive substances (TBARS) method (Ohkawa et al., 1979). 250 μl of the homogenate was mixed with 1.5 ml of each trichloroacetic acid (TCA) (20%) and TBA (0.6%). The mixture was incubated in a boiling water bath for 30 min, cooled, and 2 ml of butanol was added, mixed by vortexing, and centrifuged. The color of the butanol layer was read at 535 nm in a spectrophotometer.

## Total antioxidant capacity (TAC) in the given sample

The total antioxidant capacity is measured by the modified method of the phosphomolybdic method. (Regoli et al., 2002)

## NO scavenging activity

At physiological pH, nitric oxide generated from aqueous sodium nitroprusside solution interacts with oxygen to produce nitrite ions, which may be quantified by the Griess reaction. The activity was measured according to the Griess reaction. 1 ml of 5mM sodium nitroprusside solution was added to 100μl of the homogenized irradiated tissue sample; these reaction mixtures were incubated for 1hr at 27°Cand diluted with 1.2ml of Griess reagent (1% sulfanilamide in 5% H_3_PO_4_ and 0.1% naphthyl ethylene diaminedihydrochloride). The absorbance of the chromophore was read immediately at 550nm (Sanchez-Moreno, 2002).

### Protein Estimation

Total protein was measured according to the (Lowry et al. 1951) method keeping BSA as standard.

### DNA Damage

DNA was extracted from Wizard Genomic DNA Purification Kit, Promega, and the USA from liver and kidney tissues according to the given protocol. Isolated DNA was checked for purification at 260/280 nm ratio by spectrophotometer. Gel electrophoresis of isolated DNA was performed on 1.2% Agarose gel and visualized under UV transilluminator.

### Preparation of Sample for RNA isolation

Tissues were immediately kept in RNAlater solution, Sigma Aldrich, the USA after dissection and were kept −20° storage.

### 2.6 RNA isolation and Quantitative RT-PCR analysis

Total RNA was extracted by Trizol method (Chomczynski & Sacchi, 1987), and 2000 ng of total RNA was converted to cDNA by Prime Script RT reagent, Takara, Japan. The primer has been designed by Getprime software (https://gecftools.epfl.ch/getprime/). The primers are as mentioned in Table. 2.

**Table 2:**
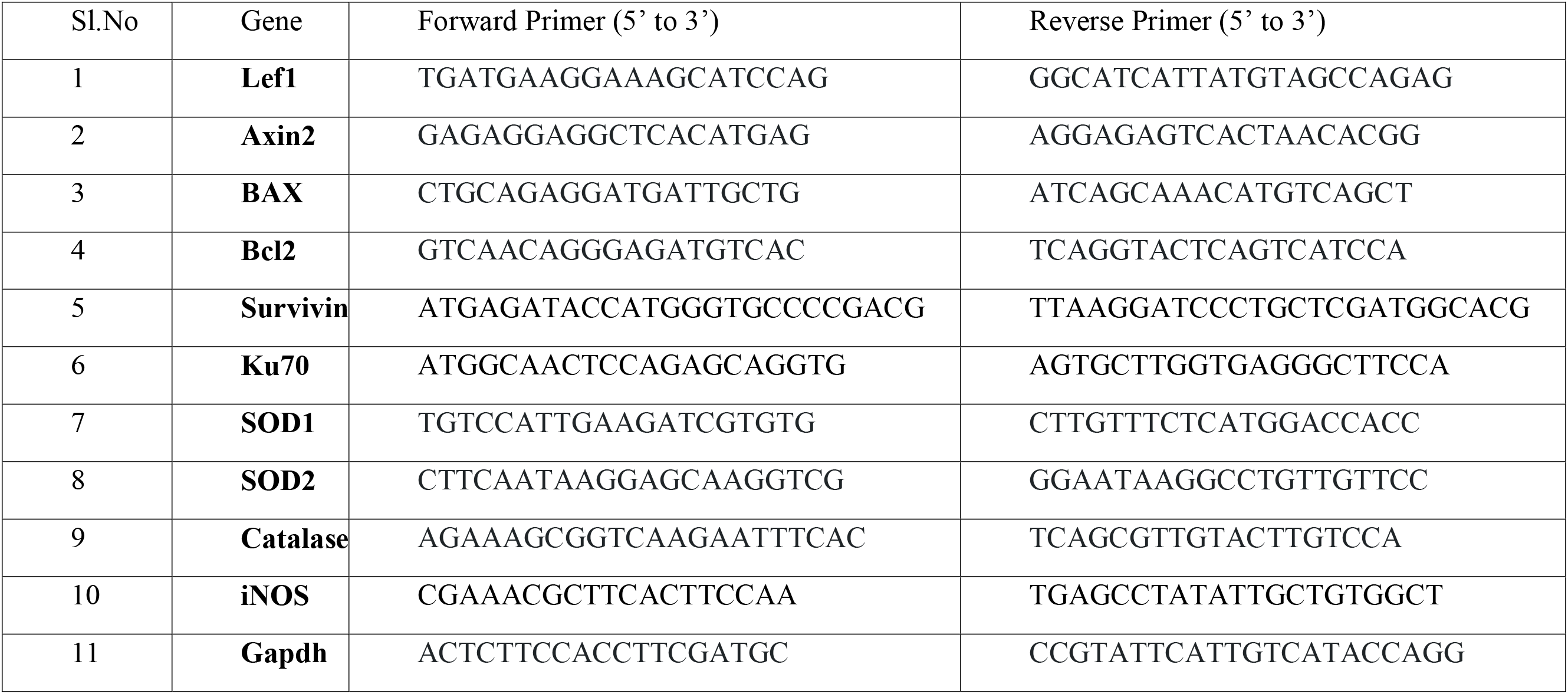
Forward and reverse primer sequence for RT-qPCR analysis:

Further cDNA was amplified with the above-given primer, and Real-Time PCR was performed on BioRad CFX-96 Real-time PCR machine. Results were analyzed by using the ΔΔCt method (Livak & Schmittgen, 2001)

### Ethical Declaration

The Institutional Animal Ethical Committee has approved the study, the University of Mysore, bearing the ethical number: UOM/IAEC/10/2016. All methods were performed under relative guidelines and regulations.

### Statistical Analysis

Results are presented as the mean ± standard error (SE) and deviation (SD) of measurements made on five mice in each group (3 in case of mRNA quantification) per experiment and is representative of separate experiments. One-way analysis of variance (ANOVA) followed by *post hoc* with Dunnett T2 was used to compare all groups and doses at all times when responses were measured at 24 hours and 30 days. Statistical differences were considered significant when *P* ≤ 0.05 using the SPSS version 22 software (Chicago) and GraphPad Prism 9.0.

## 3. Results

### 3.1 Omega 3 fish oil enhances Lef1 and Axin2 mRNA expression suggesting enhancement of the Wnt canonical pathway

In our study, there was a significant increase in the mRNA level of Lef1 and Axin2, suggesting the upregulation of the Wnt canonical pathway at post- 24 h and 30 days of irradiation when administered with Omega 3 fish oil (Figure 1 and 2, Table 3 and 4). In the liver, kidney, the relative gene expression of these genes has been increased by administrating omega 3 fish oil prior to radiation at 10Gy. Lef1 and Axin2 mRNA have significantly increased (5-60%) after irradiation compared to the control group. The administered group showed, however, a mixed recovery (up to 40%) of mRNA levels of Lef1 and Axin2, after 30 days (Figure 1 and 2, Table 3 and 4) which showed the mixed signal of the Wnt pathway following radiation exposure.

**Fig 1.**
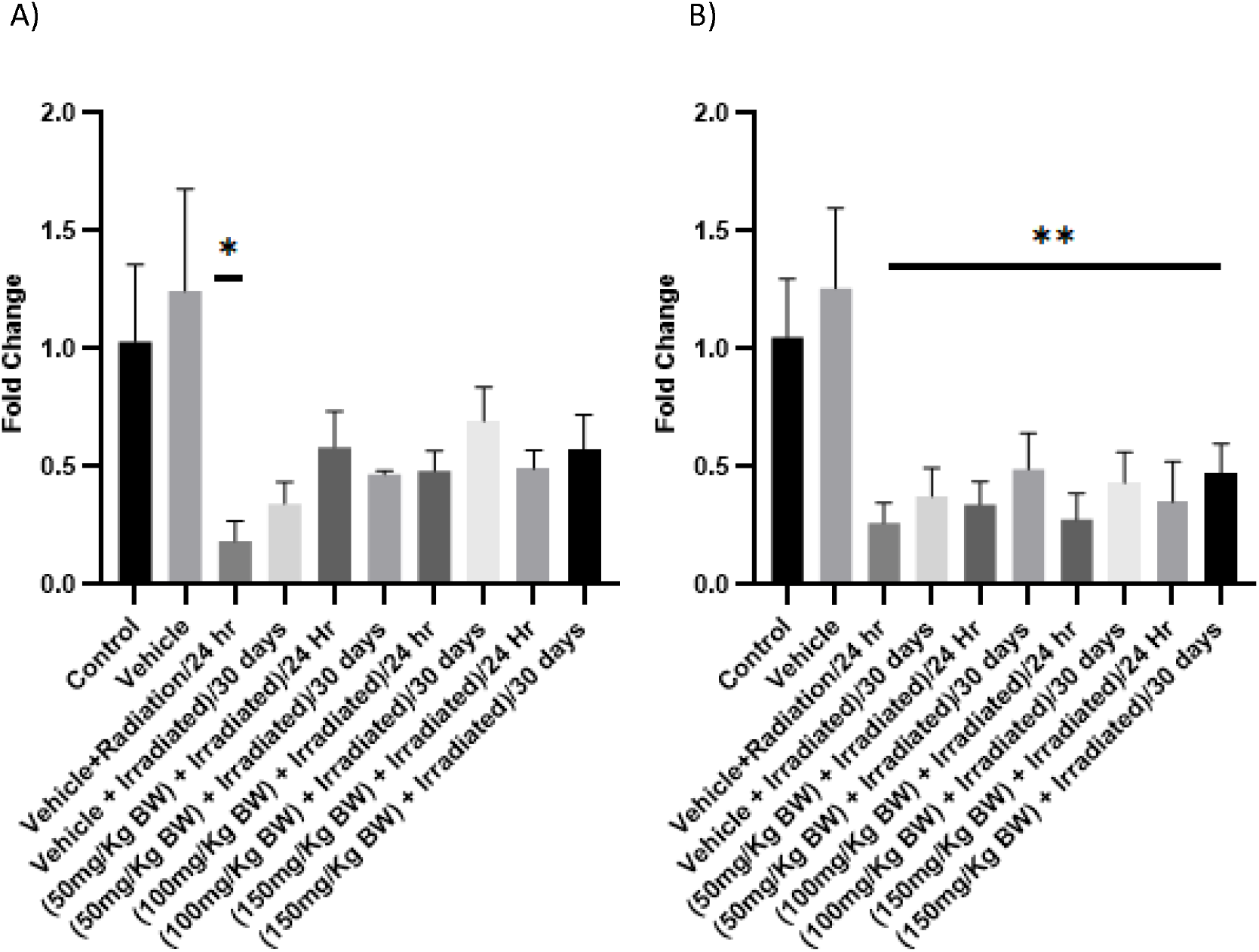
Modulatory effect of Omega 3 fatty acids on the mRNA level of Lef1 in A) Liver and B) Kidney, was measured by RT-qPCR and normalized to GAPDH mRNA (n=3; mean+ SE, one way Annova, * p<0.05, **p<0.005, ***p<0.001)

**Fig 2:**
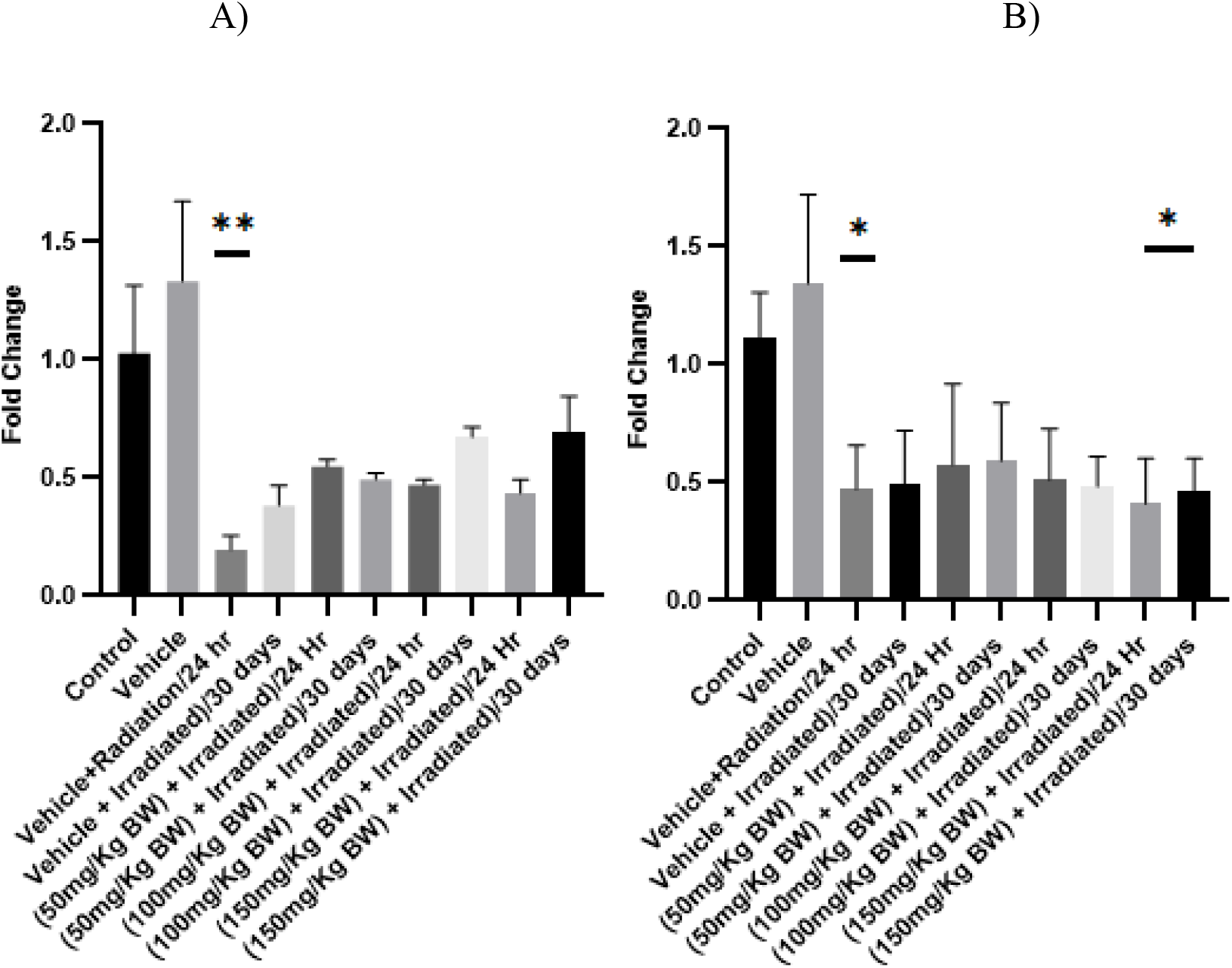
Modulatory effect of Omega 3 fatty acids on the mRNA level of Axin2 in A) Liver and B) Kidney, was measured by RT-qPCR and normalized to GAPDH mRNA (n=3; mean+ SE, one way Annova, * p<0.05, **p<0.005, ***p<0.001)

**Table 3:**
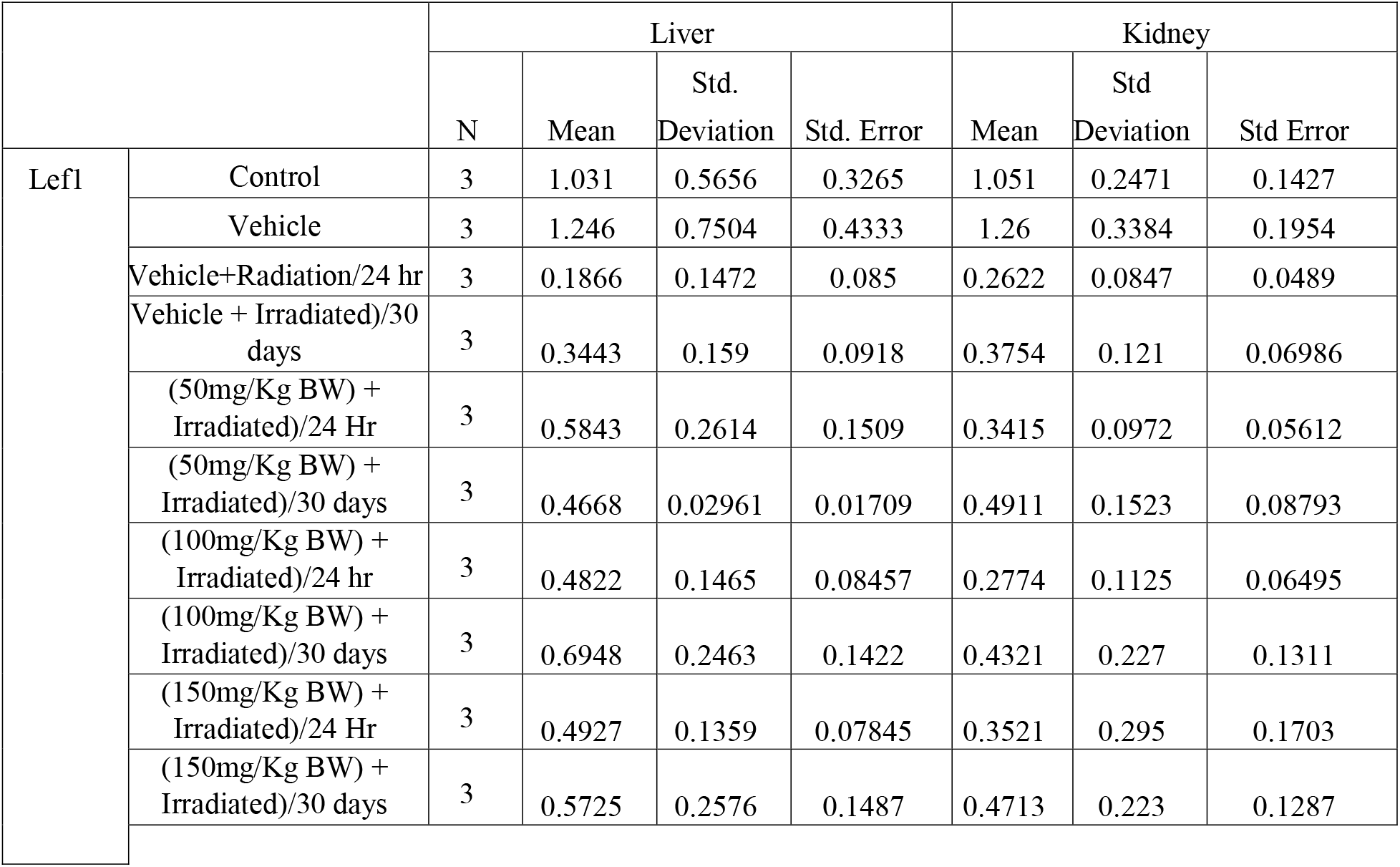
Modulatory effect of Omega 3 fatty acids on mRNA level of Lef1 expression’s descriptive value on liver, kidney.

| Lef1 |  | Liver |  |  |  | Kidney |  |  |
| --- | --- | --- | --- | --- | --- | --- | --- | --- |
|  |  | N | Mean | Std. Deviation | Std. Error | Mean | Std Deviation | Std Error |
|  | Control | 3 | 1.031 | 0.5656 | 0.3265 | 1.051 | 0.2471 | 0.1427 |
|  | Vehicle | 3 | 1.246 | 0.7504 | 0.4333 | 1.26 | 0.3384 | 0.1954 |
|  | Vehicle+Radiation/24 hr | 3 | 0.1866 | 0.1472 | 0.085 | 0.2622 | 0.0847 | 0.0489 |
|  | Vehicle + Irradiated)/30 days | 3 | 0.3443 | 0.159 | 0.0918 | 0.3754 | 0.121 | 0.06986 |
|  | (50mg/Kg BW) + Irradiated)/24 Hr | 3 | 0.5843 | 0.2614 | 0.1509 | 0.3415 | 0.0972 | 0.05612 |
|  | (50mg/Kg BW) + Irradiated)/30 days | 3 | 0.4668 | 0.02961 | 0.01709 | 0.4911 | 0.1523 | 0.08793 |
|  | (100mg/Kg BW) + Irradiated)/24 hr | 3 | 0.4822 | 0.1465 | 0.08457 | 0.2774 | 0.1125 | 0.06495 |
|  | (100mg/Kg BW) + Irradiated)/30 days | 3 | 0.6948 | 0.2463 | 0.1422 | 0.4321 | 0.227 | 0.1311 |
|  | (150mg/Kg BW) + Irradiated)/24 Hr | 3 | 0.4927 | 0.1359 | 0.07845 | 0.3521 | 0.295 | 0.1703 |
|  | (150mg/Kg BW) + Irradiated)/30 days | 3 | 0.5725 | 0.2576 | 0.1487 | 0.4713 | 0.223 | 0.1287 |

**Table 4:** Modulatory effect of Omega 3 fatty acids on mRNA level of Axin2 expression’s descriptive value on liver, kidney.

|  |  | Liver |  |  |  | Kidney |  |  |
| --- | --- | --- | --- | --- | --- | --- | --- | --- |
|  |  | N | Mean | Std. Deviation | Std. Error | Mean | Std Deviation | Std Error |
| Axin2 | Control | 3 | 1.027 | 0.5002 | 0.2888 | 1.113 | 0.1925 | 0.1111 |
|  | Vehicle | 3 | 1.335 | 0.5856 | 0.3381 | 1.345 | 0.3758 | 0.217 |
|  | Vehicle+Radiation /24 hr | 3 | 0.1955 | 0.1044 | 0.06028 | 0.4723 | 0.1856 | 0.1072 |
|  | Vehicle + Irradiated)/30 days | 3 | 0.3844 | 0.145 | 0.08374 | 0.4957 | 0.2247 | 0.1297 |
|  | (50mg/Kg BW) + Irradiated)/24 Hr | 3 | 0.5498 | 0.04702 | 0.02715 | 0.5752 | 0.3441 | 0.1987 |
|  | (50mg/Kg BW) + Irradiated)/30 days | 3 | 0.4943 | 0.0451 | 0.02604 | 0.5922 | 0.2451 | 0.1415 |
|  | (100mg/Kg BW) + Irradiated)/24 hr | 3 | 0.4708 | 0.03747 | 0.02163 | 0.5125 | 0.2175 | 0.1255 |
|  | (100mg/Kg BW) + Irradiated)/30 days | 3 | 0.6745 | 0.07123 | 0.04112 | 0.4836 | 0.2214 | 0.1278 |
|  | (150mg/Kg BW) + Irradiated)/24 Hr | 3 | 0.4361 | 0.1017 | 0.05874 | 0.4112 | 0.3324 | 0.1919 |
|  | (150mg/Kg BW) + Irradiated)/30 days | 3 | 0.6907 | 0.2681 | 0.1548 | 0.4617 | 0.244 | 0.1409 |

### 3.2 Omega 3 fish oil inhibits molecular control of apoptosis by regulating Wnt canonical pathway

The Bax and Bcl2 are candidate molecules to regulate the intrinsic apoptotic pathway. In our study, the Bax and Bcl2 in liver and kidney, is reduced when omega 3 fish oil was given. At post-24 h of irradiation (Figure 3 and 4, Table 5 and 6) the measured mRNA level was decreased in Bax and Bcl2 and the implying apoptosis was visualized in electrophoresis through an agarose gel (Figure 11) in omega 3 fish oil subjected tissues. In addition, the Wnt canonical pathway expressed survivin inhibits the activity of caspase 3 (Garg et al. 2016) resulting in the dysregulation of apoptosis and the regulation of cell survival. The mRNA level of Survivin in liver and kidney, was moderately increased in the post-24 h irradiation period in Omega 3 subjected tissues, while the fold change in tissues was found (80%) to be approximately null.

**Fig 3:**
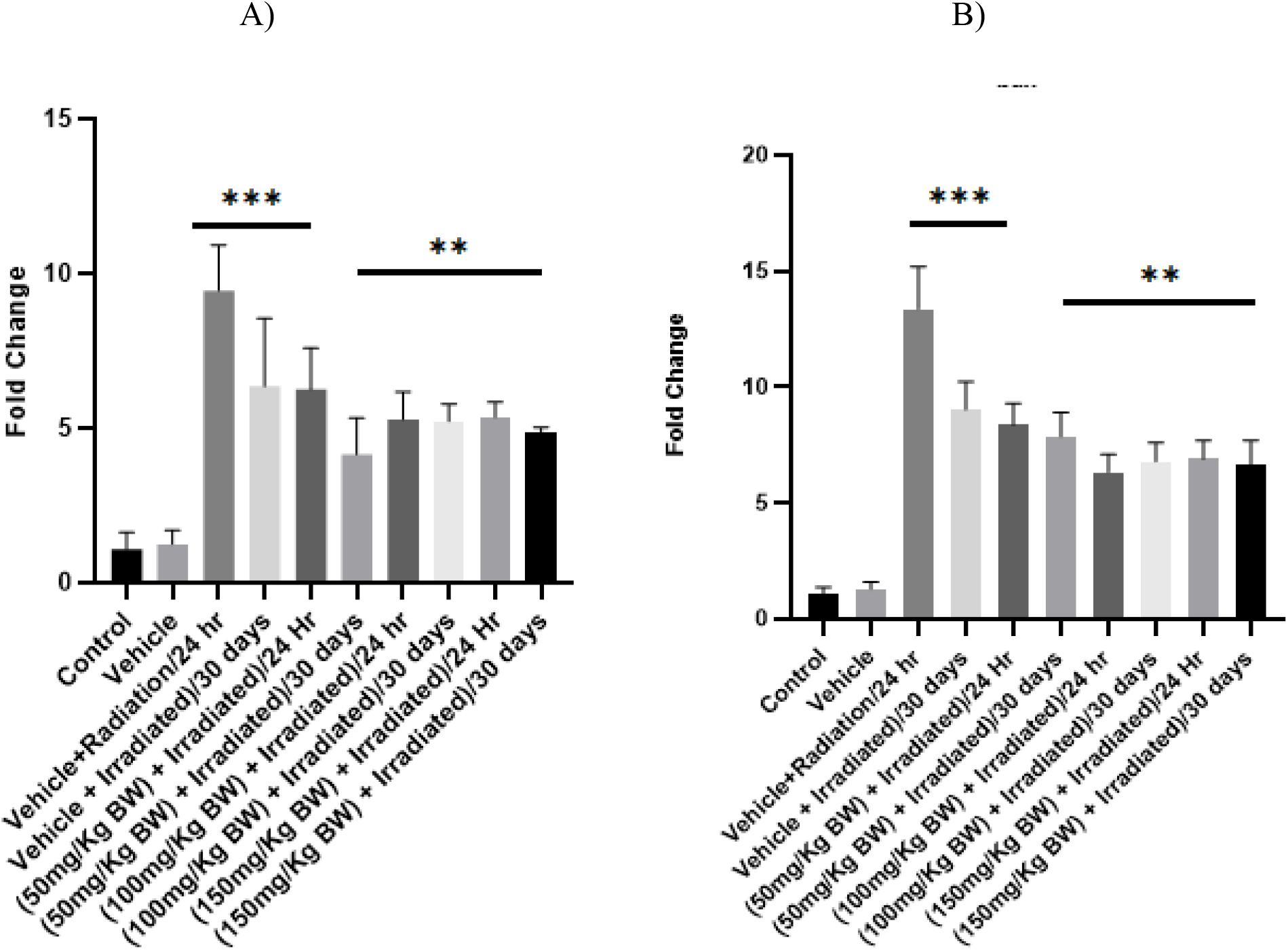
Modulatory effect of Omega 3 fatty acids on the mRNA level of Bax in A) Liver and B) Kidney, was measured by RT-qPCR and normalized to GAPDH mRNA (n=3; mean+ SE, one way Annova, * p<0.05, **p<0.005, ***p<0.001)

**Fig 4:**
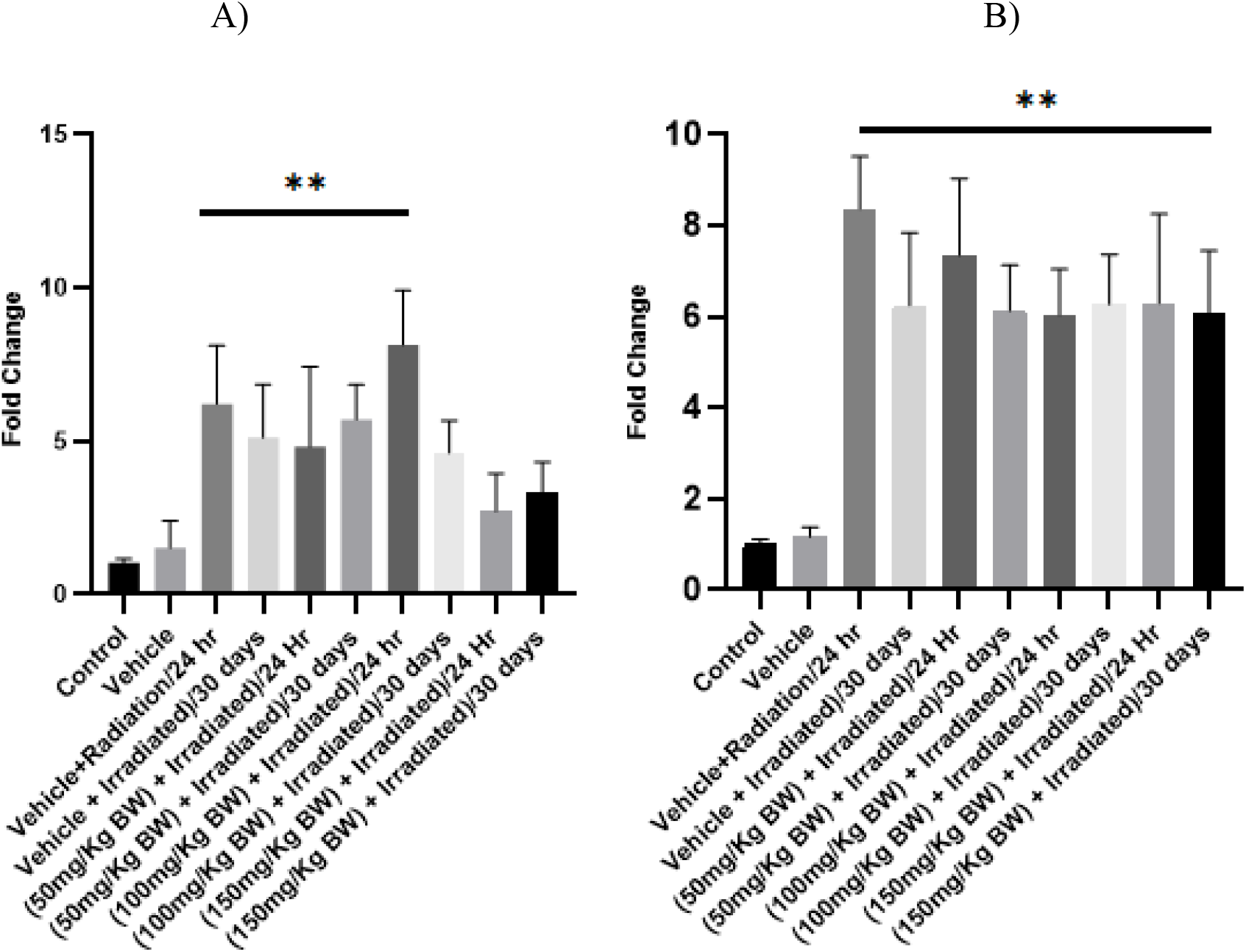
Modulatory effect of Omega 3 fatty acids on the mRNA level of Bcl2 in A) Liver and B) Kidney, was measured by RT-qPCR and normalized to GAPDH mRNA (n=3; mean+ SE, one way Annova, * p<0.05, **p<0.005, ***p<0.001)

**Fig 5.**
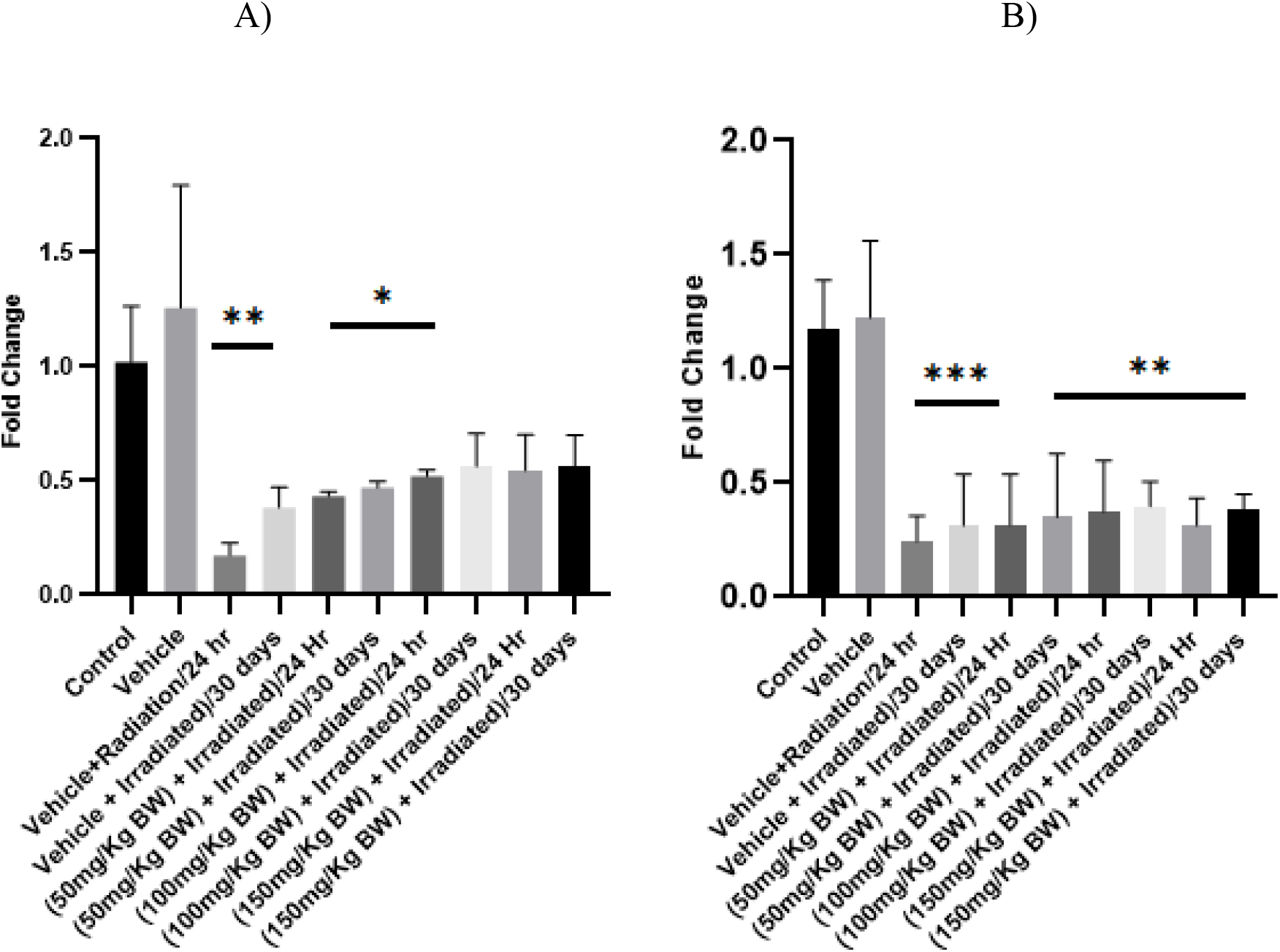
Modulatory effect of Omega 3 fatty acids on the mRNA level of Ku70 in A) Liver and B) Kidney, was measured by RT-qPCR and normalized to GAPDH mRNA (n=3; mean+ SE, one way Annova, * p<0.05, **p<0.005, ***p<0.001)

**Fig 6:**
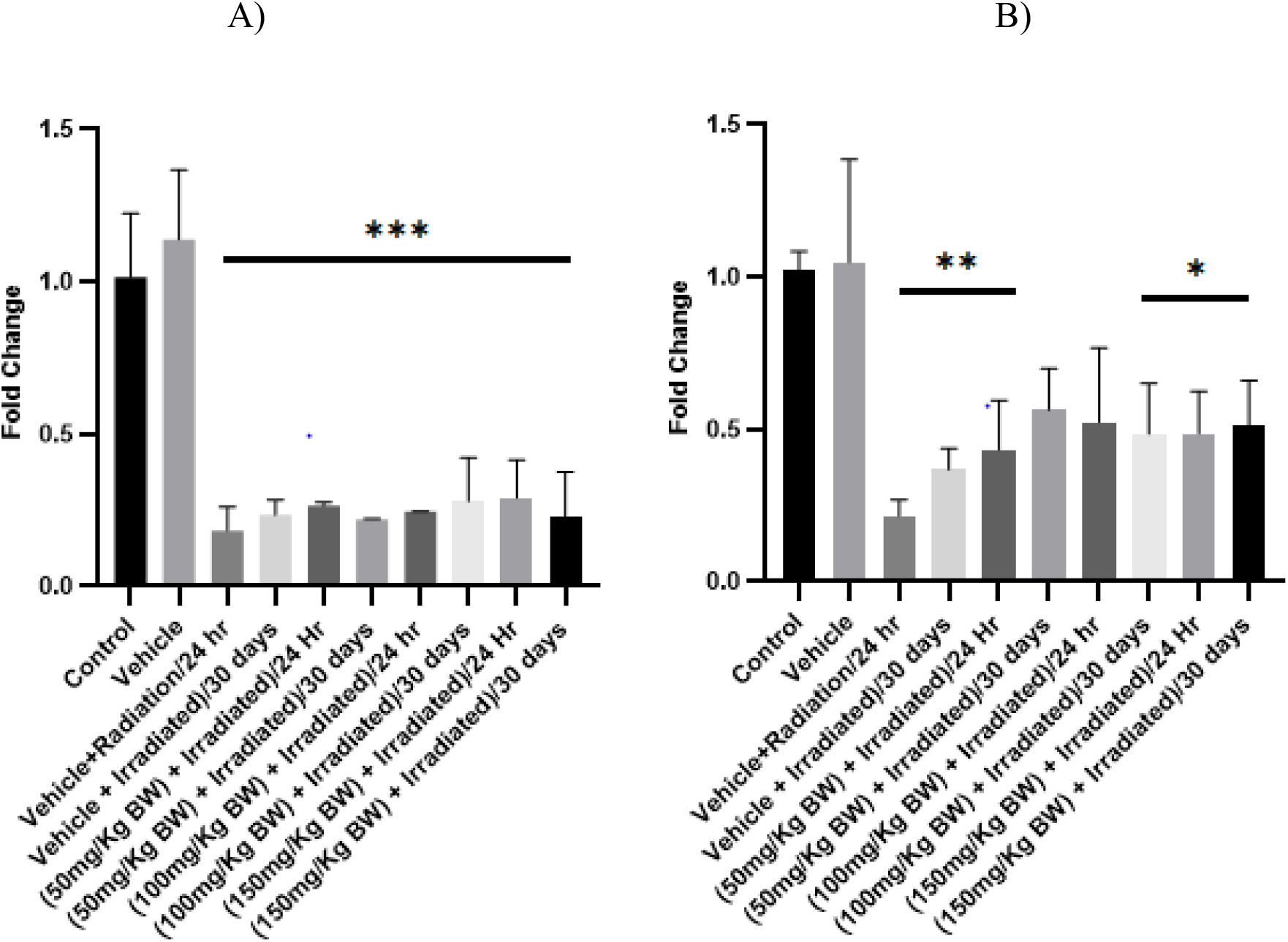
Modulatory effect of Omega 3 fatty acids on the mRNA level of Survivin in A) Liver and B) Kidney, was measured by RT-qPCR and normalized to GAPDH mRNA (n=3; mean+ SE, one way Annova, * p<0.05, **p<0.005, ***p<0.001)

**Table 5:**
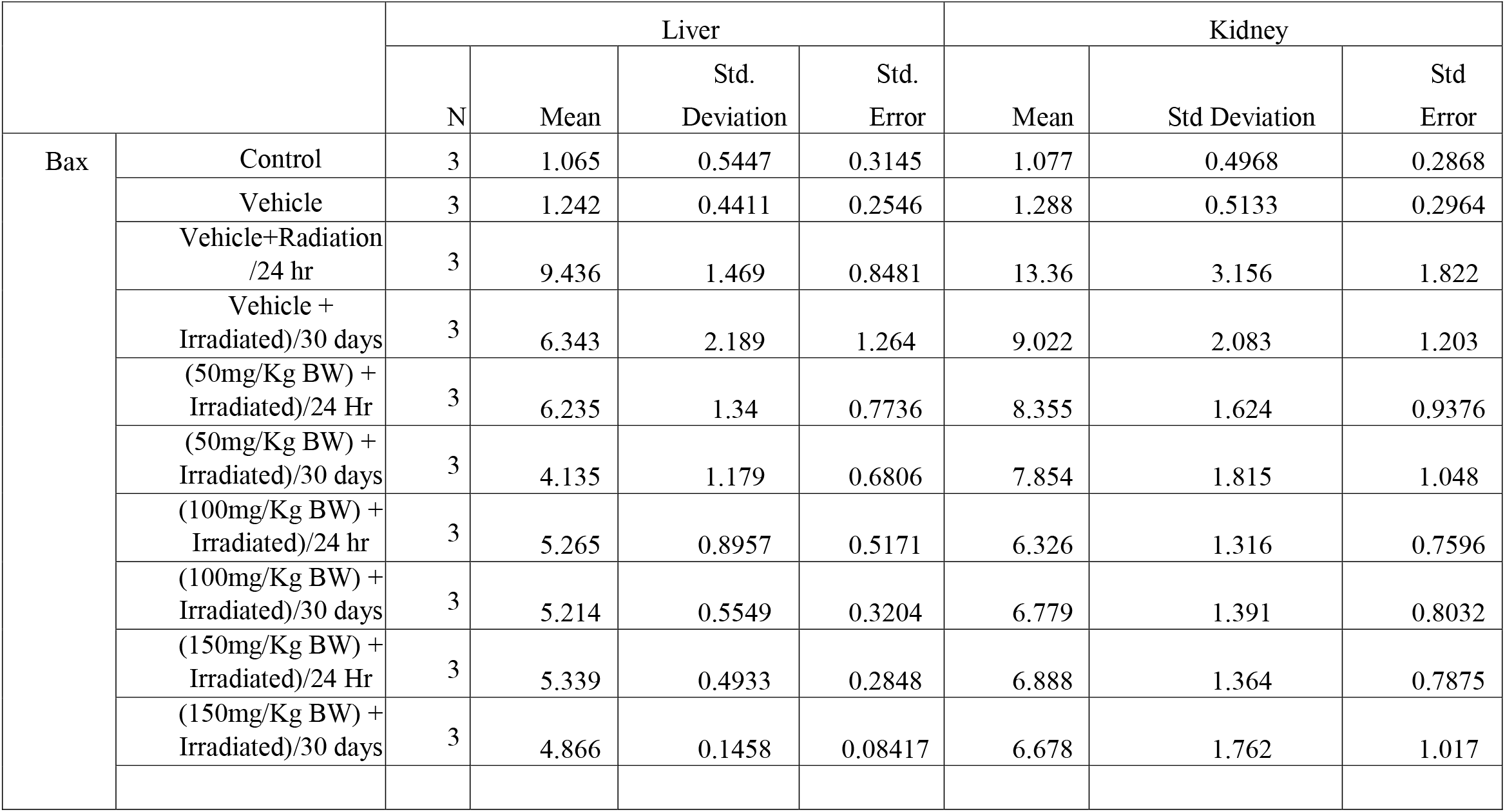
Modulatory effect of Omega 3 fatty acids on mRNA level of Bax expression’s descriptive value on liver, kidney.

**Table 6:** Modulatory effect of Omega 3 fatty acids on mRNA level of Bcl2 expression’s descriptive value on liver, kidney.

| Bcl2 |  | Liver |  |  |  | Kidney |  |  |
| --- | --- | --- | --- | --- | --- | --- | --- | --- |
|  |  | N | Mean | Std. Deviation | Std. Error | Mean | Std Deviation | Std Error |
|  | Control | 3 | 1.027 | 0.1452 | 0.0838<br>4 | 1.013 | 0.09679 | 0.05588 |
|  | Vehicle | 3 | 1.519 | 0.8836 | 0.5101 | 1.153 | 0.2234 | 0.129 |
|  | Vehicle+Radiation /24 hr | 3 | 6.225 | 1.892 | 1.092 | 8.362 | 1.156 | 0.6674 |
|  | Vehicle + Irradiated)/30 days | 3 | 5.114 | 1.738 | 1.003 | 6.242 | 1.596 | 0.9215 |
|  | (50mg/Kg BW) + Irradiated)/24 Hr | 3 | 4.823 | 2.614 | 1.509 | 7.342 | 1.683 | 0.9719 |
|  | (50mg/Kg BW) + Irradiated)/30 days | 3 | 5.71 | 1.134 | 0.655 | 6.12 | 1.005 | 0.5805 |
|  | (100mg/Kg BW) + Irradiated)/24 hr | 3 | 8.142 | 1.777 | 1.026 | 6.028 | 1.013 | 0.5846 |
|  | (100mg/Kg BW) + Irradiated)/30 days | 3 | 4.616 | 1.055 | 0.609 | 6.271 | 1.093 | 0.631 |
|  | (150mg/Kg BW) + Irradiated)/24 Hr | 3 | 2.713 | 1.225 | 0.7074 | 6.288 | 1.964 | 1.134 |
|  | (150mg/Kg BW) + Irradiated)/30 days | 3 | 3.308 | 1.013 | 0.5846 | 6.077 | 2.367 | 1.367 |

### 3.3 Omega 3 fish oil decreases Molecular control of radiotoxicity coupled with the Wnt canonical pathway via oxidative stress

The mRNA level of SOD1 and SOD2 was significantly reduced (30–90%), indicating the effect on the Superoxide dismutase enzyme after 24 h and 30 days of irradiation. Administration of Omega 3 fish oil enhances SOD1 and SOD2 expression to some extent, showing recovery up to 40-60% (Figure 7 and 8, Table 9 and 10). The SOD scavenging activity was also increased after omega 3 fish oil was administered at 24-h and 30 days post irradiation (Figure 12; Table 13). During the post-radiation period of 30 days, the mRNA level of the liver, kidney, SOD1 and SOD2 were improved. Further, the administration of omega 3 fish oil slightly improved the mRNA level of catalase in the liver, kidneys (Figure 9, Table 11). In the post-radiation period of 30 days, the Catalase level has been slightly increased after administration of omega 3 fish oil (Figure 14, Table 15). Similarly, the iNOS level of significantly improved in omega 3 administered mice except for germ cells (Figure 10 Table 12). Mice showed improved SOD activity when treated with omega-3 fish oil. 30 days of post-irradiation reveals a mild recovery of SOD activity, but still lower than control (Figure 12; Table 13). Catalase shows recovery in the liver and kidney in omega-3-administered mice (Figure 14; Table 15). Even after the omega 3 administration, the elevated level of LPO (Figure 13; Table 14) was observed in all conditions.

**Fig 7:**
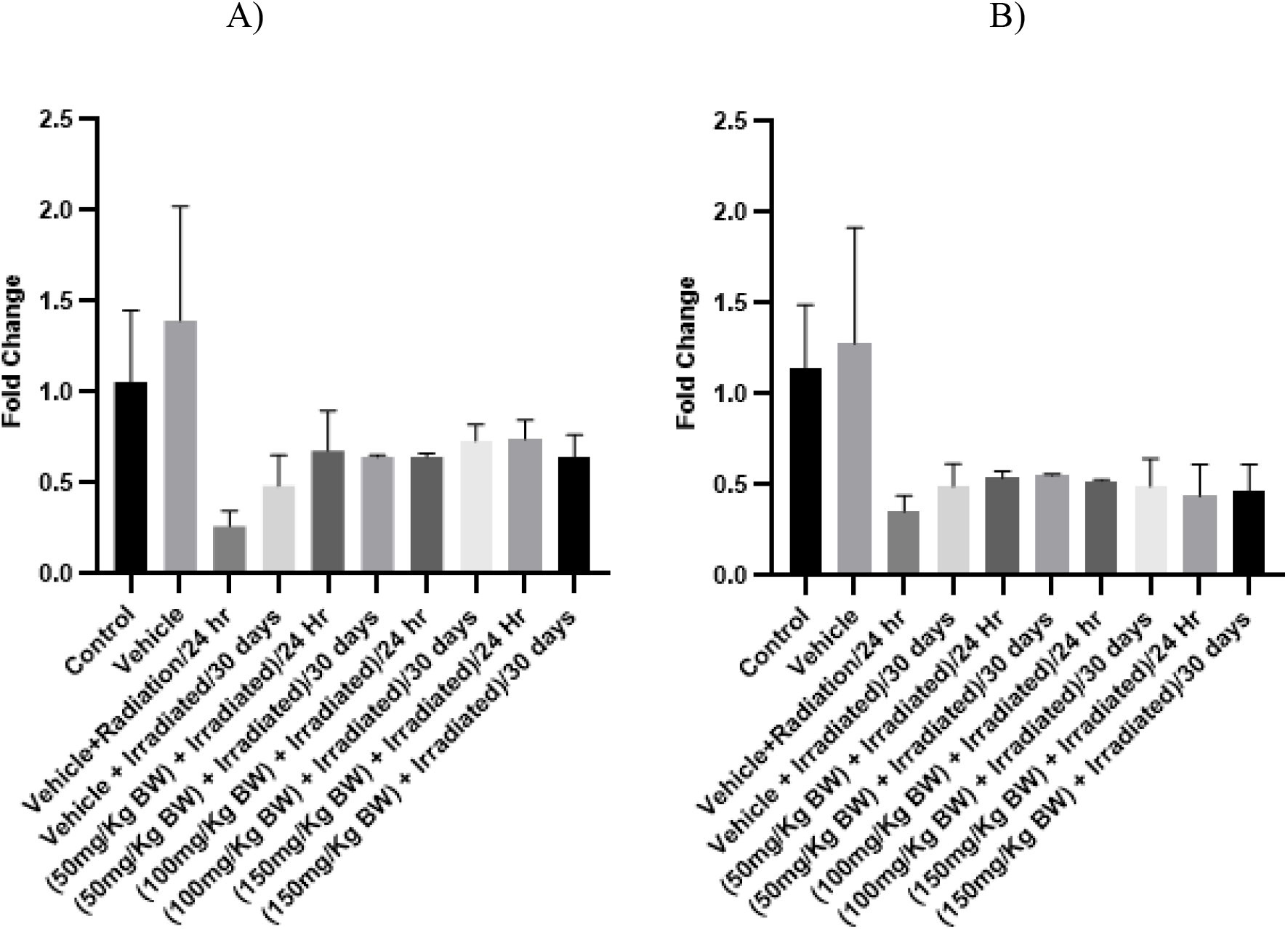
Modulatory effect of Omega 3 fatty acids on the mRNA level of SOD1 in A) Liver and B) Kidney, was measured by RT-qPCR and normalized to GAPDH mRNA (n=3; mean+ SE, one way Annova, * p<0.05, **p<0.005, ***p<0.001)

**Fig 8:**
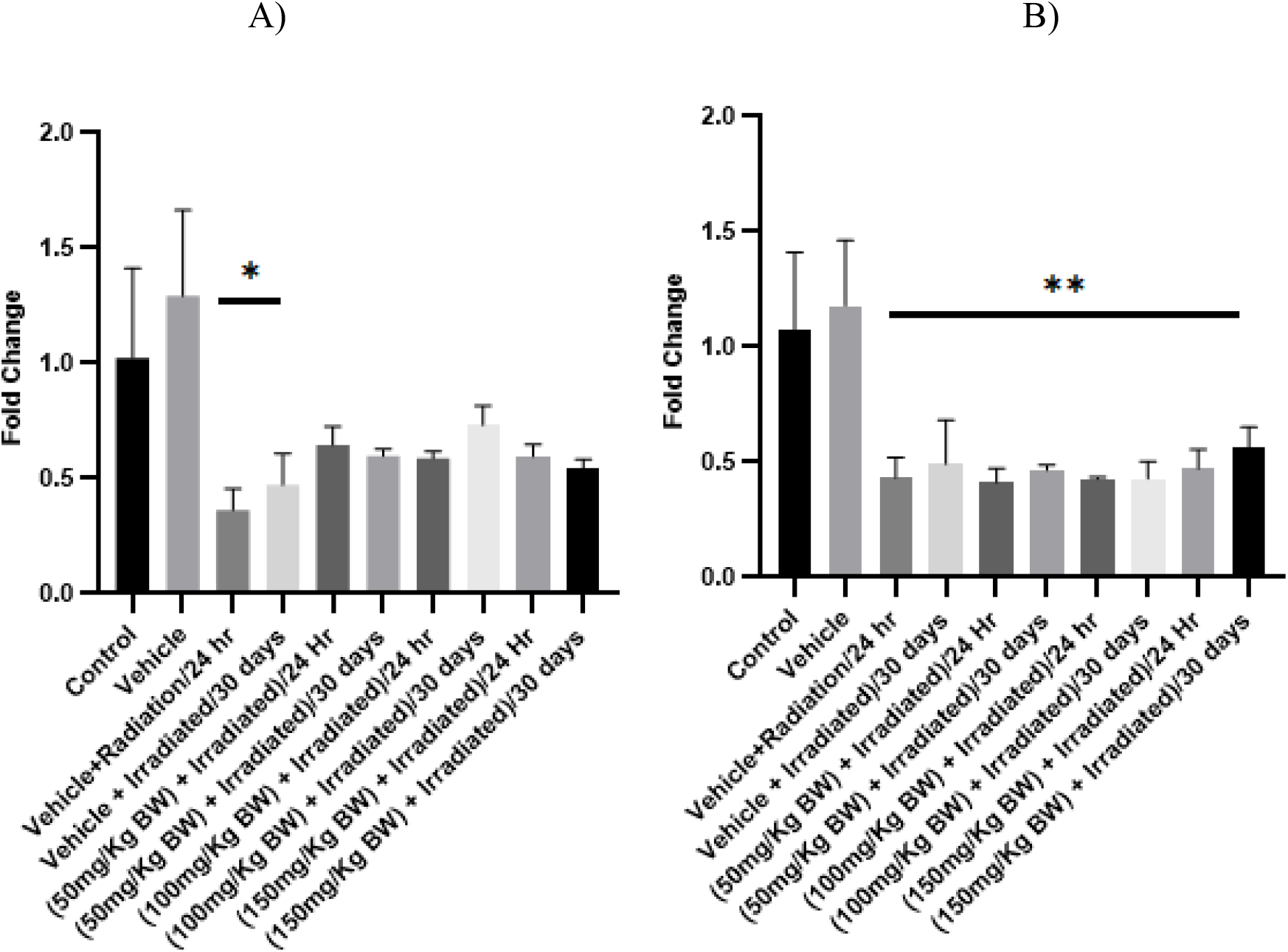
Modulatory effect of Omega 3 fatty acids on the mRNA level of SOD2 in A) Liver and B) Kidney, was measured by RT-qPCR and normalized to GAPDH mRNA (n=3; mean+ SE, one way Annova, * p<0.05, **p<0.005, ***p<0.001)

**Fig 9.**
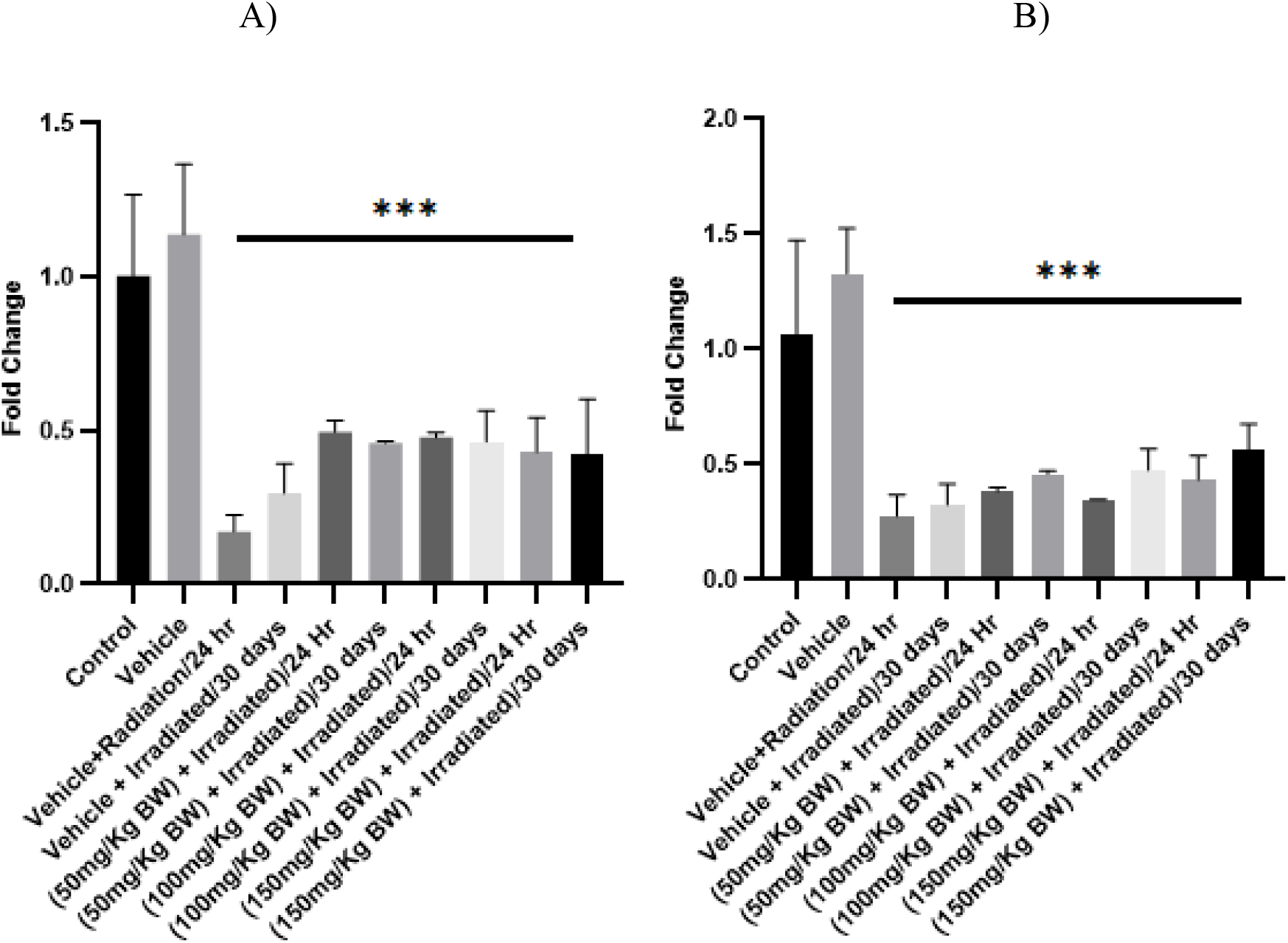
Modulatory effect of Omega 3 fatty acids on the mRNA level of Catalase in A) Liver and B) Kidney, was measured by RT-qPCR and normalized to GAPDH mRNA (n=3; mean+ SE, one way Annova, * p<0.05, **p<0.005, ***p<0.001)

**Fig 10.**
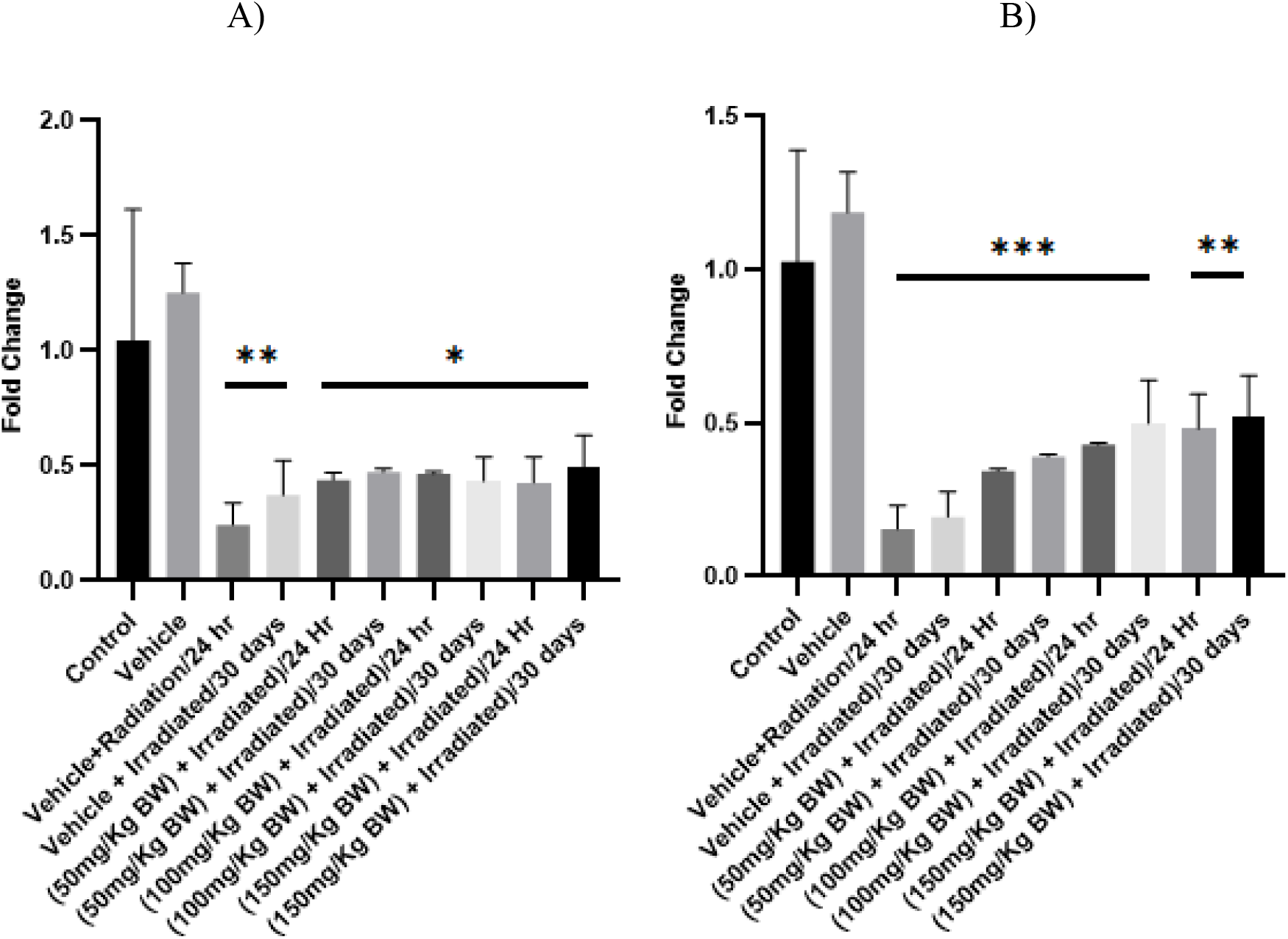
Modulatory effect of Omega 3 fatty acids on the mRNA level of iNOS in A) Liver and B) Kidney, was measured by RT-qPCR and normalized to GAPDH mRNA (n=3; mean+ SE, one way Annova, * p<0.05, **p<0.005, ***p<0.001)

**Fig 11.**
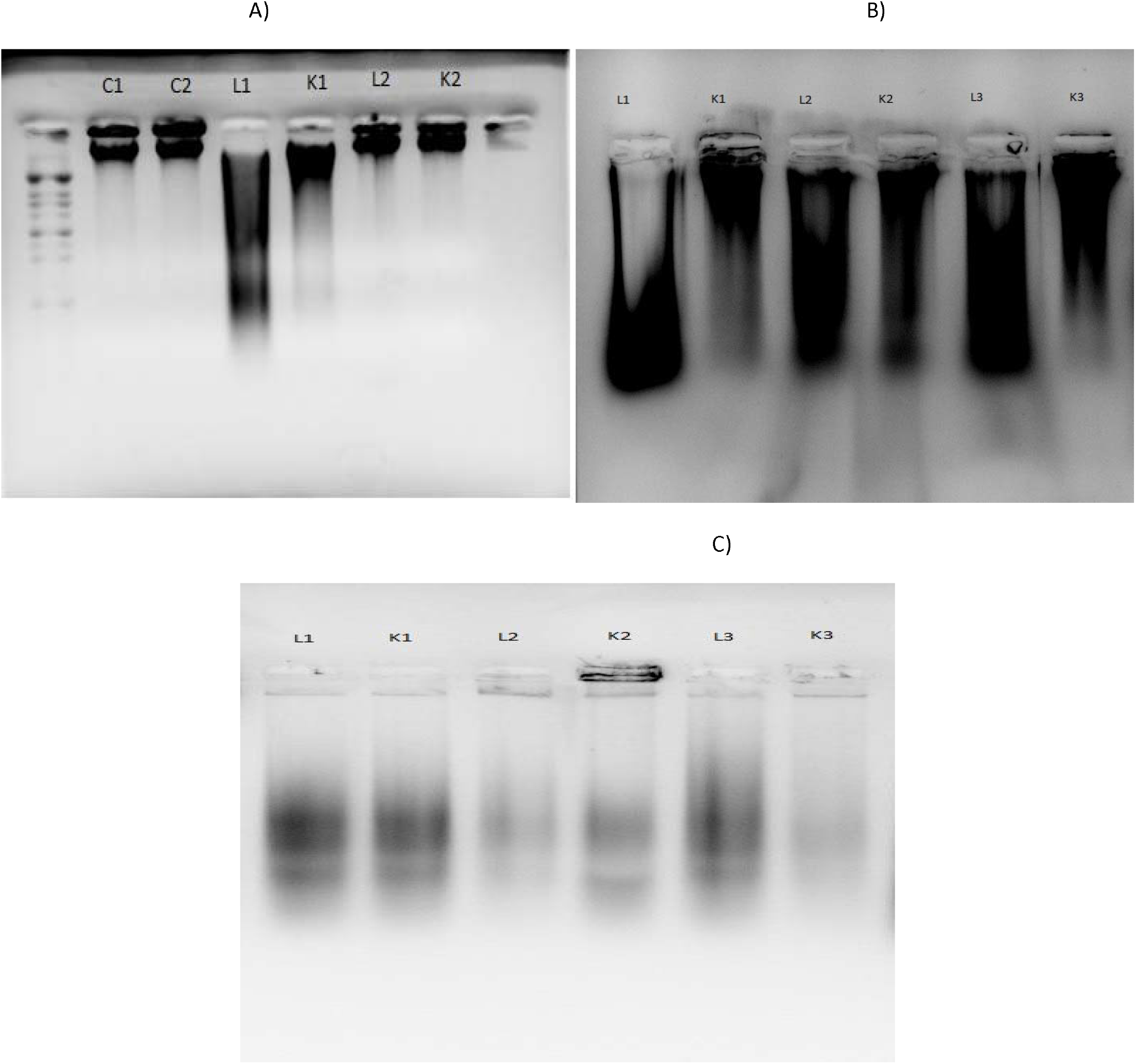
Agarose gel electrophoresis of A) Control liver (C1) and kidney(C2), Post 24hr irradiation with 10Gy liver (L1) and kidney (K1), Post 30 days of irradiation with 10Gy liver (L2) and kidney (K2), B) Agarose gel electrophoresis of 24hr Post irradiated 10Gy with supplementation of 50mg/kg BW Omega 3 fish oil in liver (L1) and kidney (K1), 100mg/kg BW Omega 3 fish oil in liver (L2) and kidney (K2), 150mg/kg BW Omega 3 fish oil in liver (L3) and kidney (K3) C) Agarose gel electrophoresis of 30 days Post irradiated 10Gy with supplementation of 50mg/kg BW Omega 3 fish oil in liver (L1) and kidney (K1), 100mg/kg BW Omega 3 fish oil in liver (L2) and kidney (K2), 150mg/kg BW Omega 3 fish oil in liver (L3) and kidney (K3).

**Fig 12.**
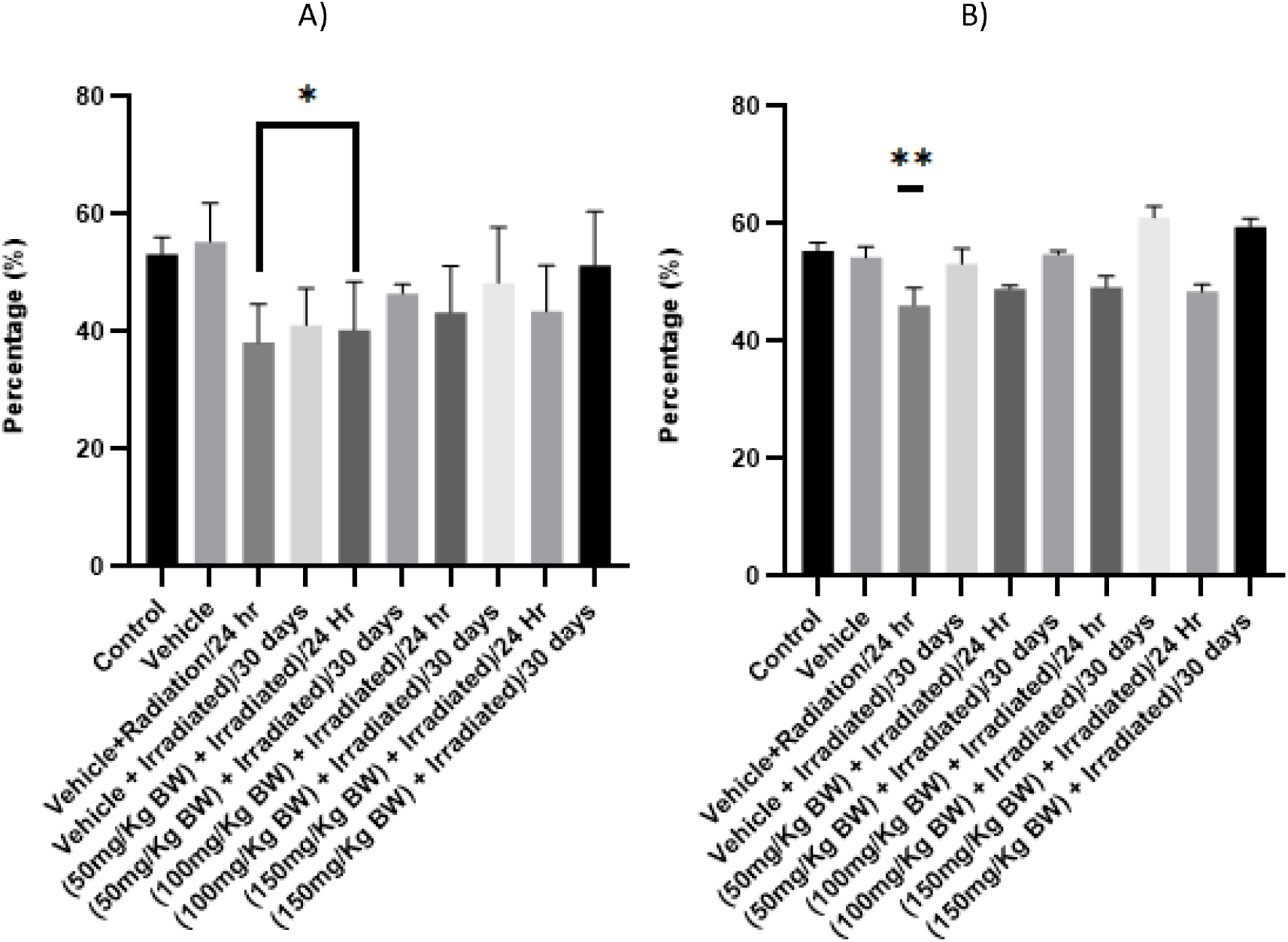
Modulatory effect of Omega 3 fatty acids on the SOD scavenging activity in A) Liver, B) Kidney (n=5; mean ± SE, one way annova, * p<0.05, **p<0.005, ***p<0.001

**Fig 13.**
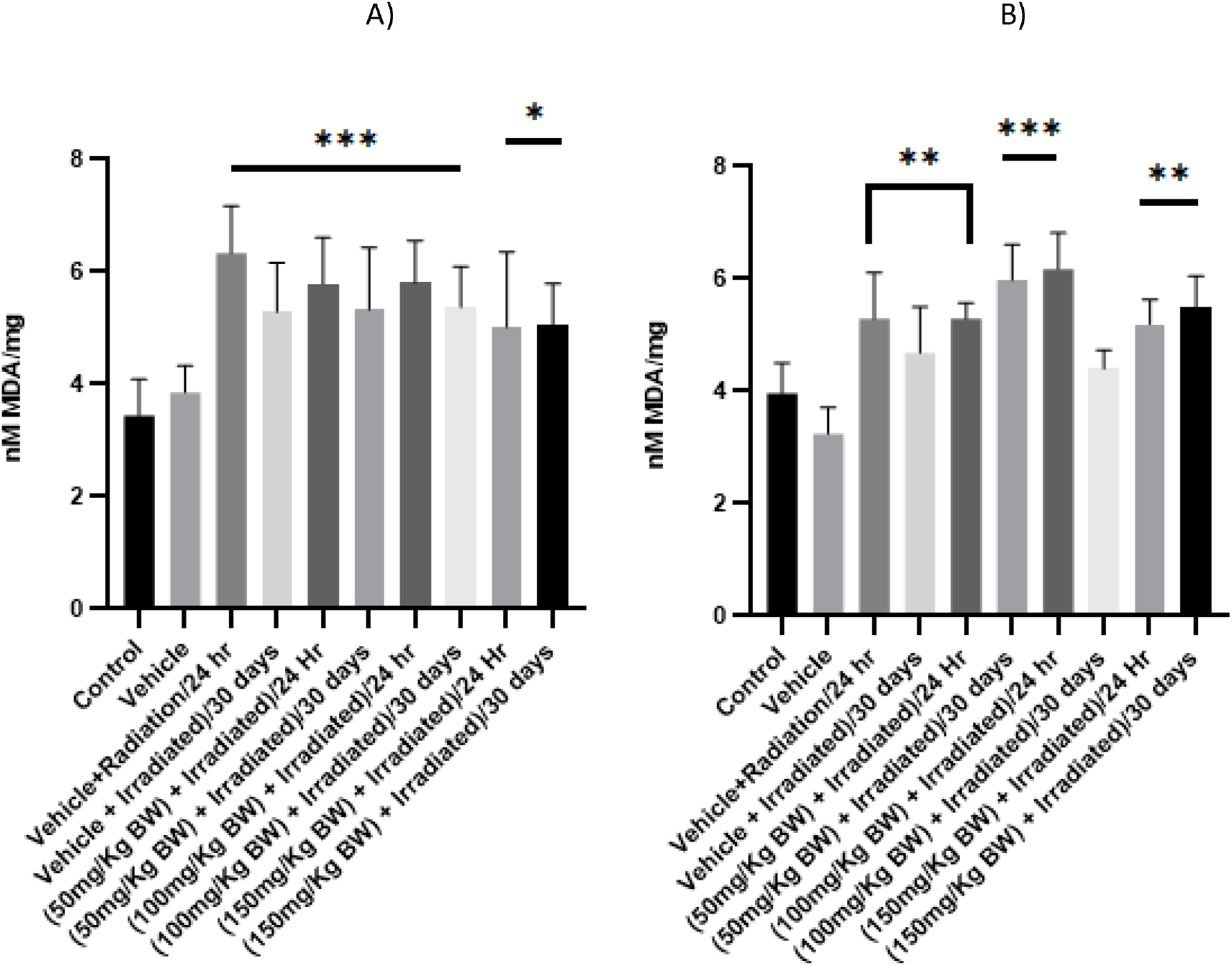
Modulatory effect of Omega 3 fatty acids on the Lipid Peroxidation in A) Liver, B) Kidney (n=5; mean ± SE, one way annova, * p<0.05, **p<0.005, ***p<0.001

**Fig 14.**
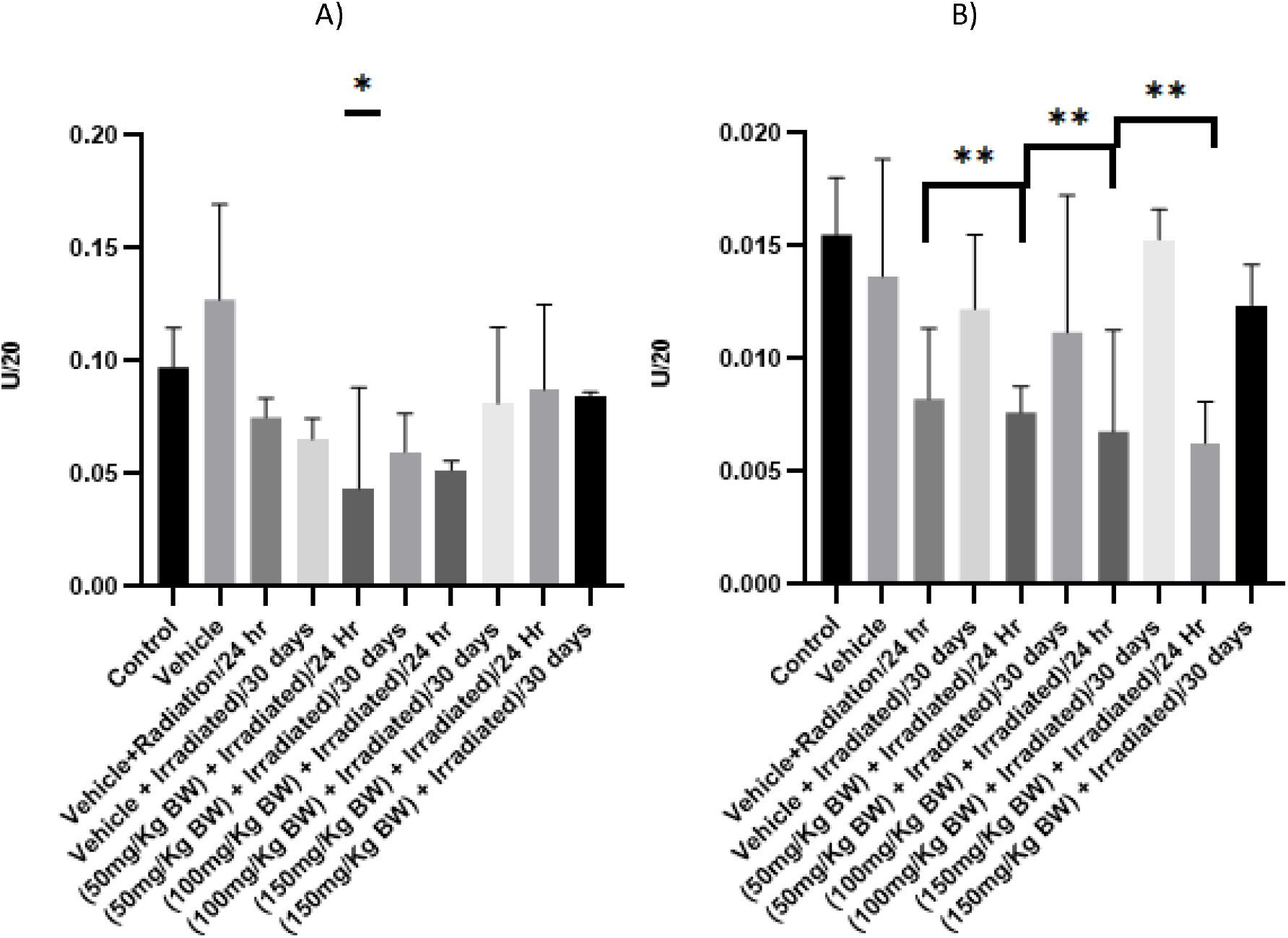
Modulatory effect of Omega 3 fatty acids on the Catalase activity in A) Liver, B) Kidney (n=5; mean ± SE, one way annova, * p<0.05, **p<0.005, ***p<0.001

**Fig 15.**
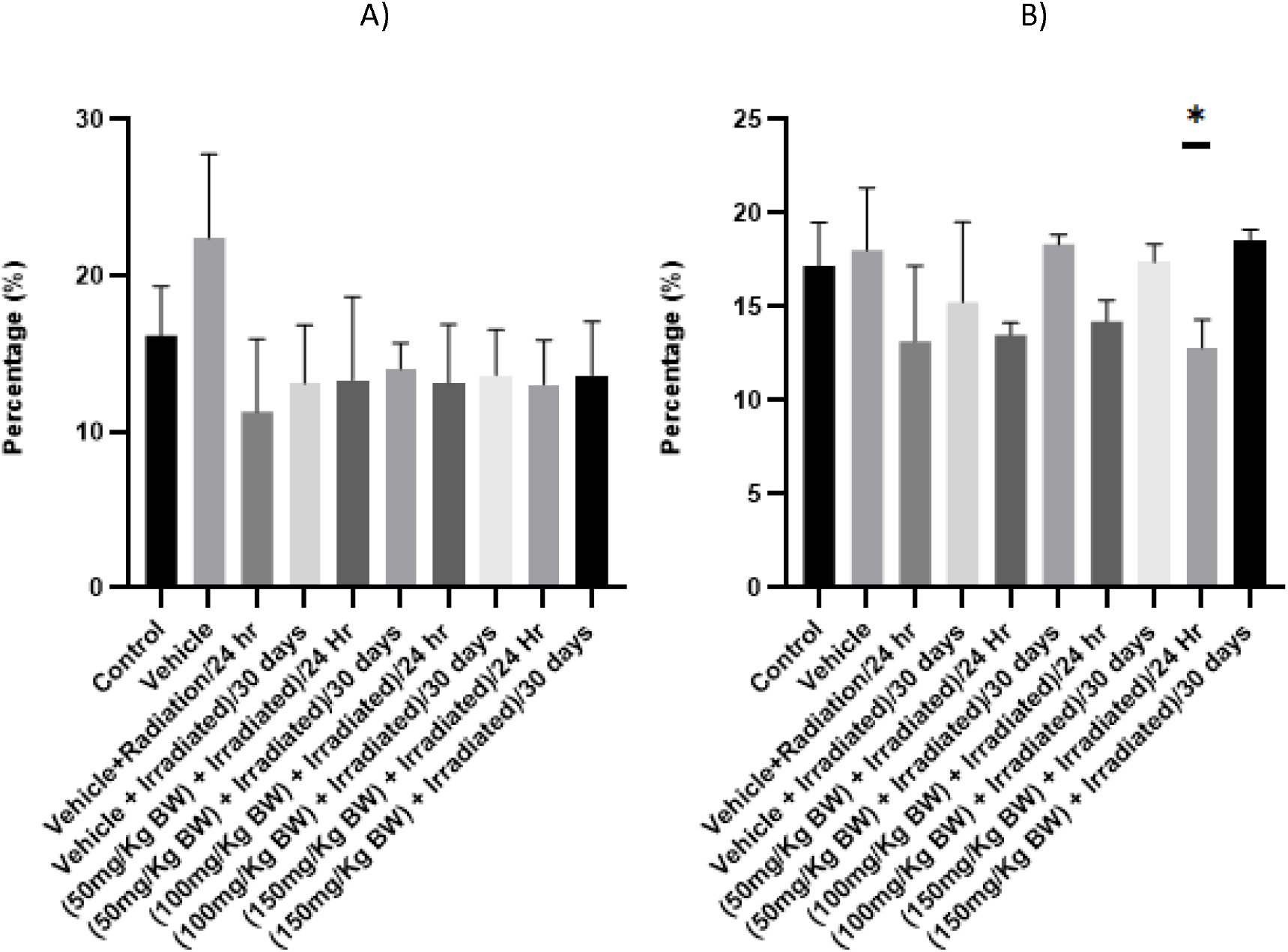
Modulatory effect of Omega 3 fatty acids on the NO scavenging activity in A) Liver, B) Kidney (n=5; mean ± SE, one way annova, * p<0.05, **p<0.005, ***p<0.001

**Table 7:**
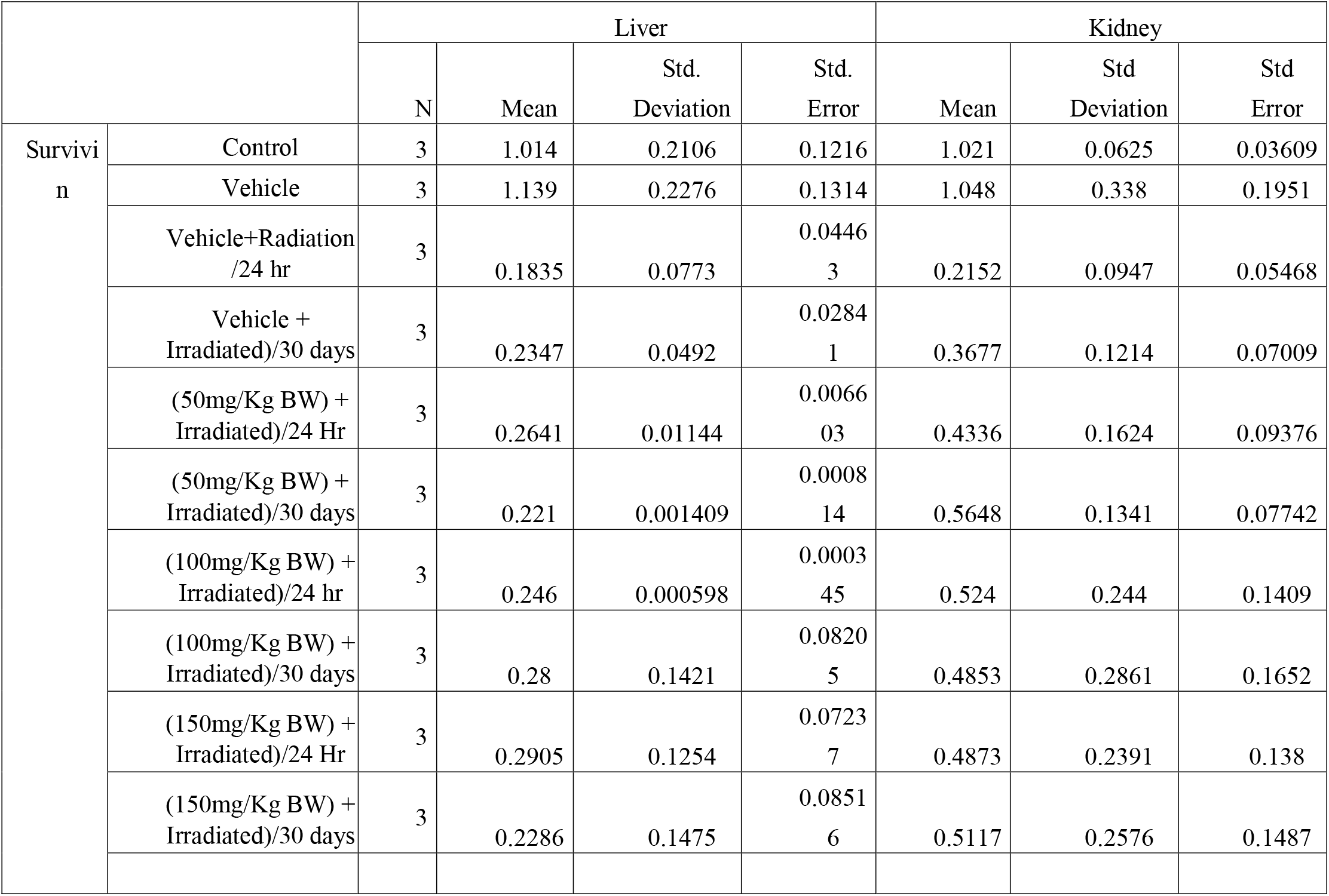
Modulatory effect of Omega 3 fatty acids on mRNA level of Survivin expression’s descriptive value on liver, kidney.

**Table 8:**
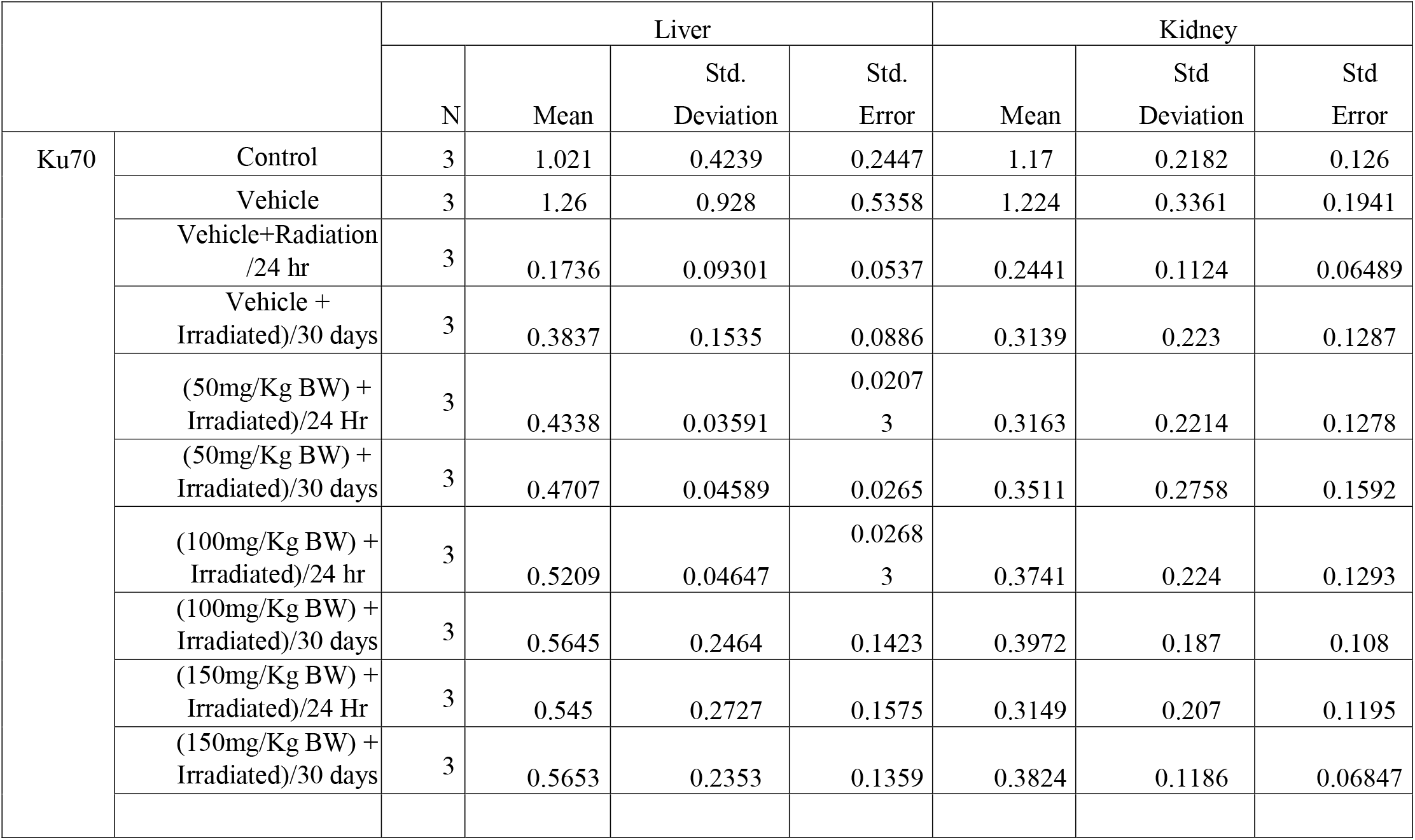
Modulatory effect of Omega 3 fatty acids on mRNA level of Ku70 expression’s descriptive value on liver, kidney, spleen.

**Table 9:** Modulatory effect of Omega 3 fatty acids on mRNA level of SOD1 expression’s descriptive value on liver, kidney, spleen.

| SOD1 |  | N | Liver |  |  | Kidney |  |  |
| --- | --- | --- | --- | --- | --- | --- | --- | --- |
|  |  |  | Mean | Std. Deviation | Std. Error | Mean | Std Deviation | Std Error |
|  | Control | 3 | 1.057 | 0.6826 | 0.3941 | 1.141 | 0.6082 | 0.3511 |
|  | Vehicle | 3 | 1.395 | 1.087 | 0.6277 | 1.278 | 1.105 | 0.6382 |
|  | Vehicle+Radiation /24 hr | 3 | 0.2642 | 0.1425 | 0.0823 | 0.3526 | 0.1529 | 0.08825 |
|  | Vehicle + Irradiated)/30 days | 3 | 0.4845 | 0.2927 | 0.169 | 0.4929 | 0.2111 | 0.1219 |
|  | (50mg/Kg BW) + Irradiated)/24 Hr | 3 | 0.6788 | 0.3825 | 0.2209 | 0.5384 | 0.06682 | 0.03858 |
|  | (50mg/Kg BW) + Irradiated)/30 days | 3 | 0.6447 | 0.01337 | 0.007721 | 0.554 | 0.01229 | 0.007093 |
|  | (100mg/Kg BW) + Irradiated)/24 hr | 3 | 0.6455 | 0.0314 | 0.01813 | 0.523 | 0.01317 | 0.007606 |
|  | (100mg/Kg BW) + Irradiated)/30 days | 3 | 0.7314 | 0.1583 | 0.09142 | 0.493 | 0.264 | 0.1524 |
|  | (150mg/Kg BW) + Irradiated)/24 Hr | 3 | 0.7424 | 0.1814 | 0.1047 | 0.4378 | 0.3043 | 0.1757 |
|  | (150mg/Kg BW) + Irradiated)/30 days | 3 | 0.6421 | 0.215 | 0.1242 | 0.4669 | 0.2535 | 0.1464 |

**Table 10:**
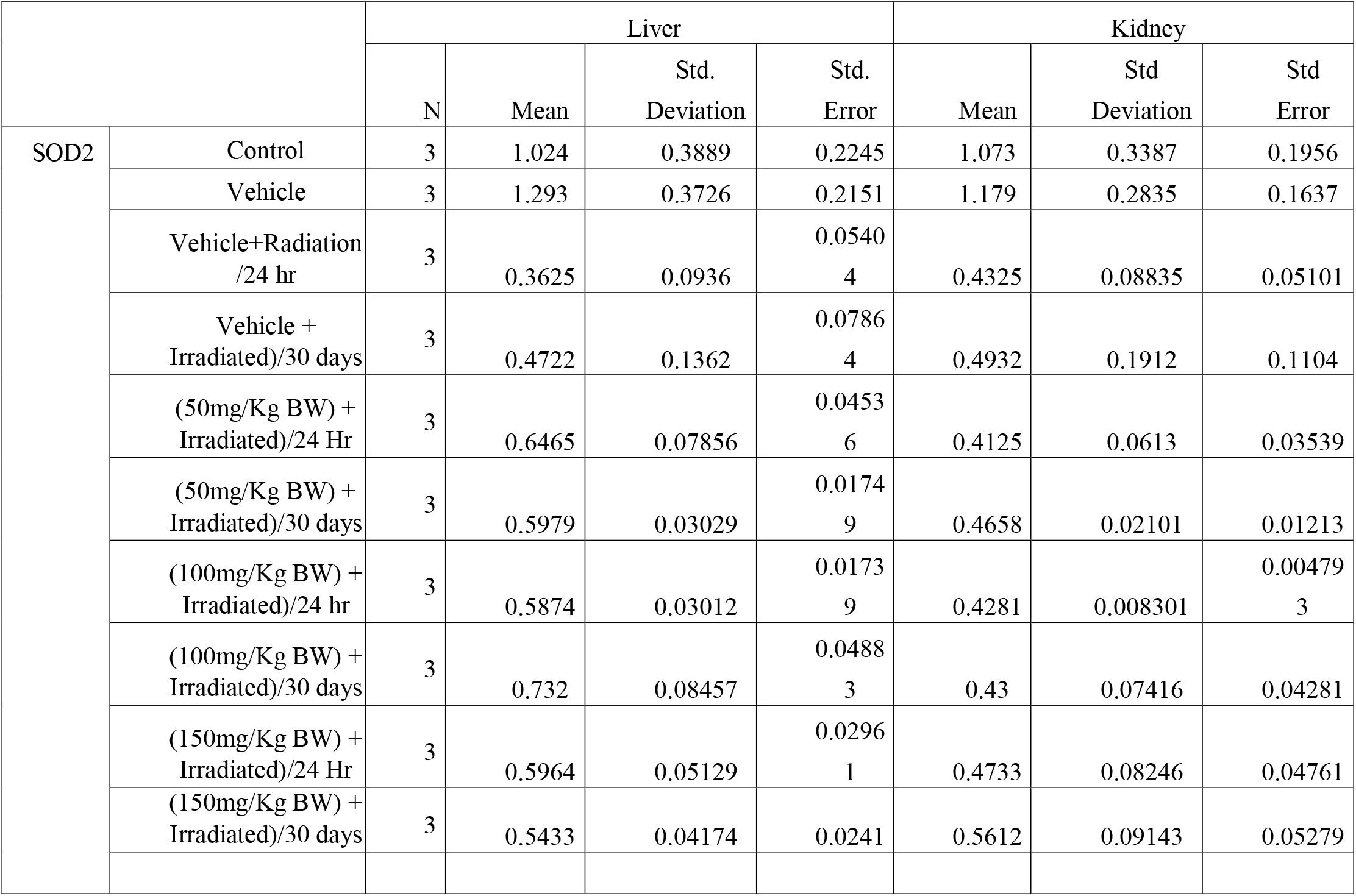
Modulatory effect of Omega 3 fatty acids on mRNA level of SOD2 expression’s descriptive value on liver, kidney.

**Table 11:** Modulatory effect of Omega 3 fatty acids on mRNA level of Catalase expression’s descriptive value on liver, kidney, spleen.

|  |  | Liver |  |  |  | Kidney |  |  |
| --- | --- | --- | --- | --- | --- | --- | --- | --- |
|  |  | N | Mean | Std. Deviation | Std. Error | Mean | Std Deviation | Std Error |
| Cat | Control | 3 | 1.005 | 0.455 | 0.2627 | 1.064 | 1.578 | 0.9109 |
|  | Vehicle | 3 | 1.138 | 0.3959 | 0.2286 | 1.329 | 0.3425 | 0.1977 |
|  | Vehicle+Radiation /24 hr | 3 | 0.1723 | 0.09284 | 0.0536 | 0.2756 | 0.1625 | 0.0938 |
|  | Vehicle + Irradiated)/30 days | 3 | 0.2973 | 0.1669 | 0.0963<br>5 | 0.3264 | 0.1536 | 0.08867 |
|  | (50mg/Kg BW) + Irradiated)/24 Hr | 3 | 0.4953 | 0.06595 | 0.0380<br>8 | 0.3847 | 0.02765 | 0.01596 |
|  | (50mg/Kg BW) + Irradiated)/30 days | 3 | 0.4612 | 0.00768 | 0.0044<br>36 | 0.4596 | 0.0231 | 0.01334 |
|  | (100mg/Kg BW) + Irradiated)/24 hr | 3 | 0.4779 | 0.02793 | 0.0161<br>2 | 0.3484 | 0.00224 | 0.001295 |
|  | (100mg/Kg BW) + Irradiated)/30 days | 3 | 0.4646 | 0.1757 | 0.1014 | 0.4765 | 0.1583 | 0.09142 |
|  | (150mg/Kg BW) + Irradiated)/24 Hr | 3 | 0.4305 | 0.1948 | 0.1125 | 0.4342 | 0.1788 | 0.1033 |
|  | (150mg/Kg BW) + Irradiated)/30 days | 3 | 0.4249 | 0.3091 | 0.1785 | 0.5647 | 0.1911 | 0.1103 |

**Table 12:**
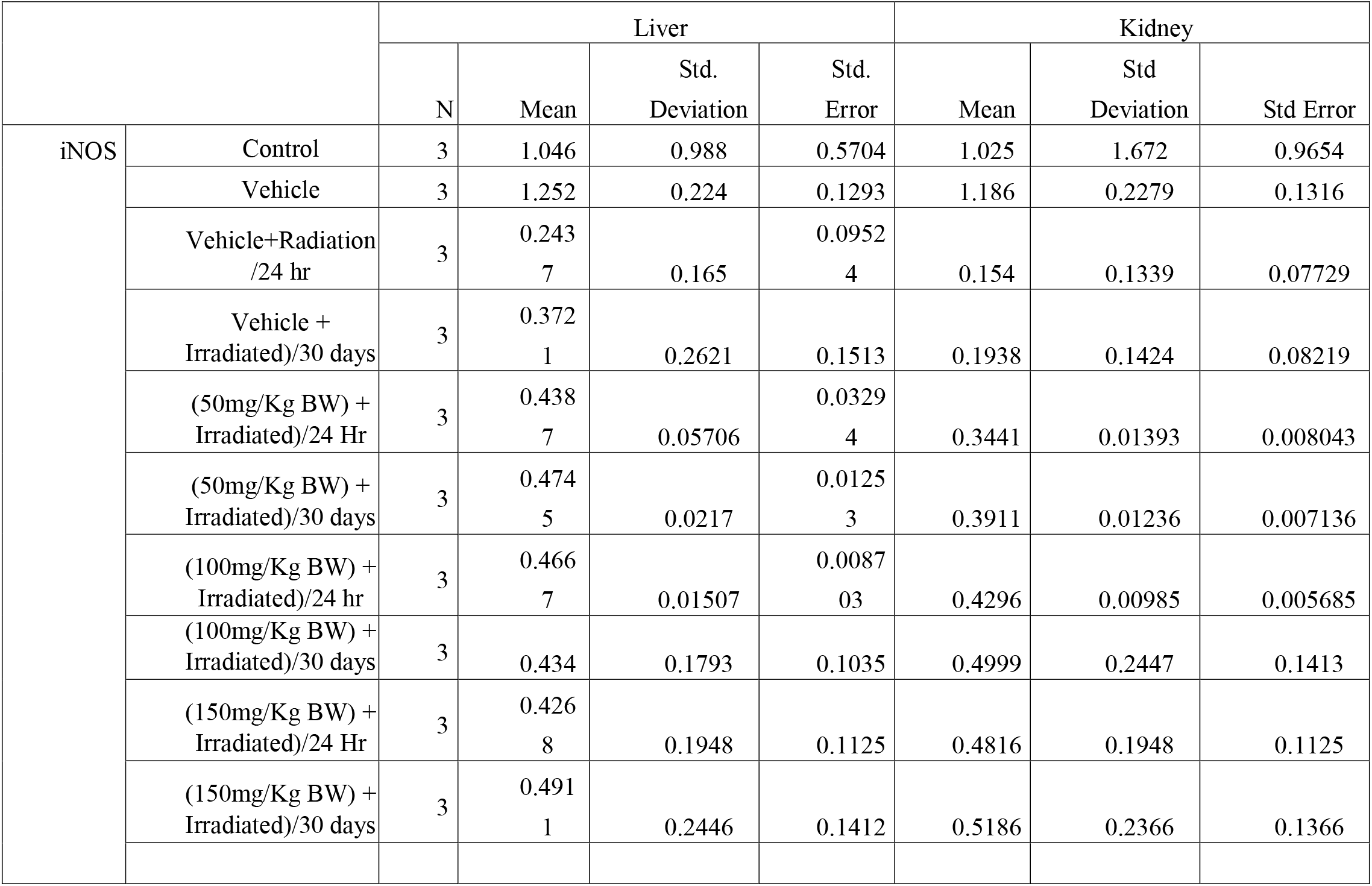
Modulatory effect of Omega 3 fatty acids on mRNA level of iNOS expression’s descriptive value on liver, kidney, spleen.

**Table 13:** Modulatory effect of Omega 3 fatty acids on Superoxide dismutase scavenging activity (%) on radiation induced radiotoxicity in liver, kidney.

|  | Liver |  | Kidney |  |
| --- | --- | --- | --- | --- |
|  | Mean | Standard deviation | Mean | Standard deviation |
| Control | 53.24 | 2.958947951 | 55.45 | 3.12627574 |
| Vehicle | 55.3746 | 6.5173 | 54.3257 | 4.1108 |
| Vehicle+Radiation/24 hr | 38.2427 | 6.4725 | 46.1857 | 6.5071 |
| Vehicle + Irradiated)/30 days | 41.14438 | 6.2812 | 53.247125 | 5.7875 |
| (50mg/Kg BW) + Irradiated)/24 Hr | 40.3575 | 8.212047046 | 48.955 | 1.148520788 |
| (50mg/Kg BW) + Irradiated)/30 days | 46.57 | 1.554627501 | 54.82 | 1.317042141 |
| (100mg/Kg BW) + Irradiated)/24 hr | 43.365 | 7.890472856 | 49.225 | 4.136121372 |
| (100mg/Kg BW) + Irradiated)/30 days | 48.345 | 9.508824681 | 61.0675 | 4.334600135 |
| (150mg/Kg BW) + Irradiated)/24 Hr | 43.4825 | 7.834622705 | 48.5275 | 2.610177197 |
| (150mg/Kg BW) + Irradiated)/30 days | 51.31 | 9.214232909 | 59.5675 | 3.069890063 |

### 3.4 Omega 3 fish oil suppress lipid peroxidation and enhance Total antioxidant capacity in radiation induced membrane oxidation

Our study shown elevated level of lipid peroxidation and suppression in Total antioxidant capacity (Fig 13 and 16; Table: 14 and 17) when exposed to radiation alone but it has shown recovery when added with omega 3 fish oil. Omega 3 fish oil also enhances recovery after 30 days of recovery period. TAC has been improved when treated with omega 3 fish oil and kept for 30 days recovery period.

**Fig 16.**
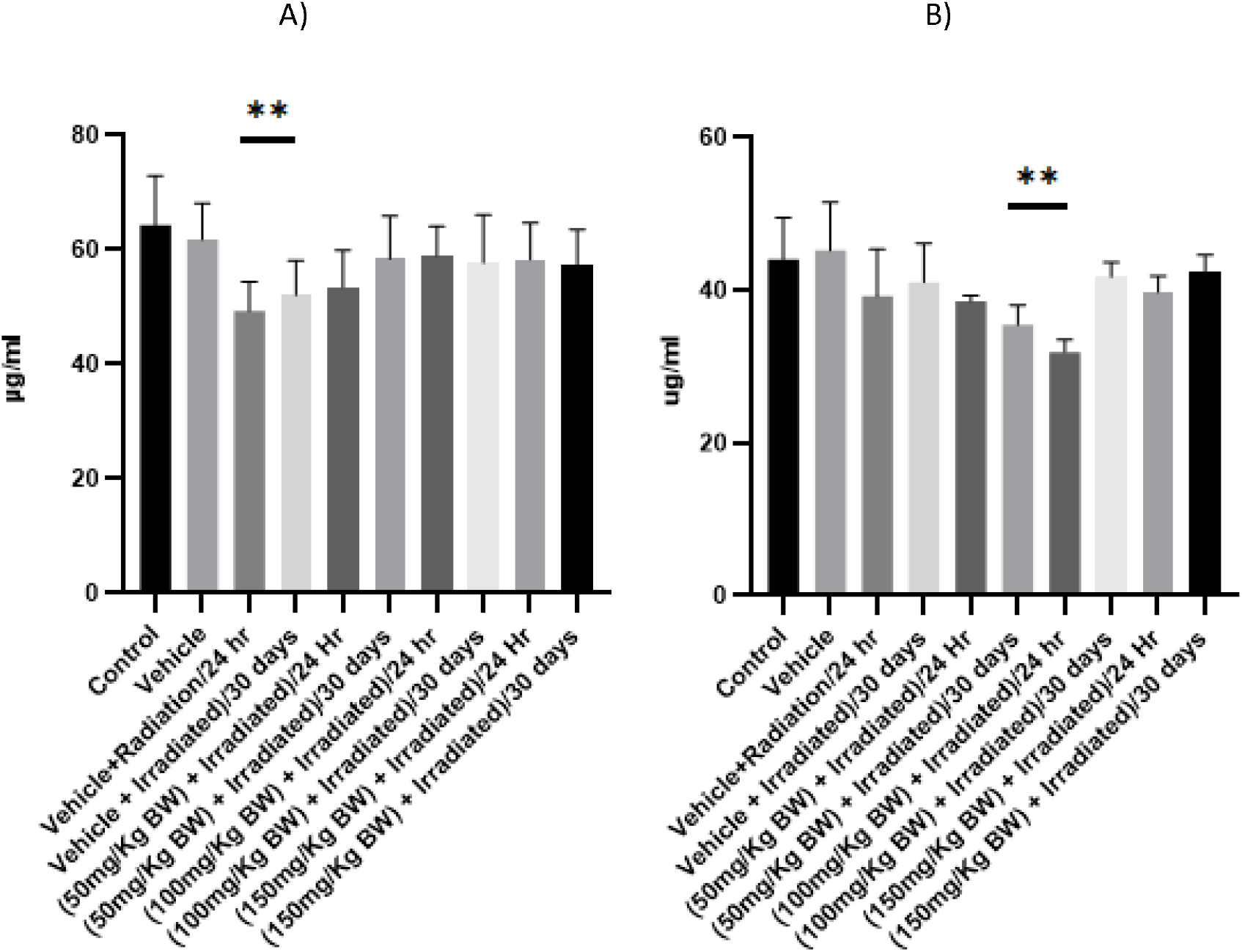
Modulatory effect of Omega 3 fatty acids on the Total Antioxidant Capacity in A) Liver, B) Kidney (n=5; mean ± SE, one way annova, * p<0.05, **p<0.005, ***p<0.001

**Table 14:** Modulatory effect of Omega 3 fatty acids on lipid peroxidation (nM) of radiation induced radiotoxicity in liver, kidney.

|  | Liver |  | Kidney |  |
| --- | --- | --- | --- | --- |
|  | Mean | Standard Deviation | Mean | Standard deviation |
| Control | 3.441 | 0.646912186 | 3.96 | 1.215 |
| Vehicle | 3.8571 | 0.47385 | 3.246 | 1.034 |
| Vehicle+Radiation/24 hr | 6.3281 | 0.8472 | 5.29 | 1.838 |
| Vehicle + Irradiated)/30 days | 5.28917 | 0.8735 | 4.682 | 1.836 |
| (50mg/Kg BW) + Irradiated)/24 Hr | 5.7925 | 0.820088134 | 5.293 | 0.6304 |
| (50mg/Kg BW) + Irradiated)/30 days | 5.3275 | 1.1046983 | 5.985 | 1.393 |
| (100mg/Kg BW) + Irradiated)/24 hr | 5.81 | 0.74418987 | 6.175 | 1.451 |
| (100mg/Kg BW) + Irradiated)/30 days | 5.38 | 0.717199094 | 4.4 | 0.7552 |
| (150mg/Kg BW) + Irradiated)/24 Hr | 5.015 | 1.338682744 | 5.198 | 0.9634 |
| (150mg/Kg BW) + Irradiated)/30 days | 5.0625 | 0.729781949 | 5.485 | 1.258 |

**Table 15:**
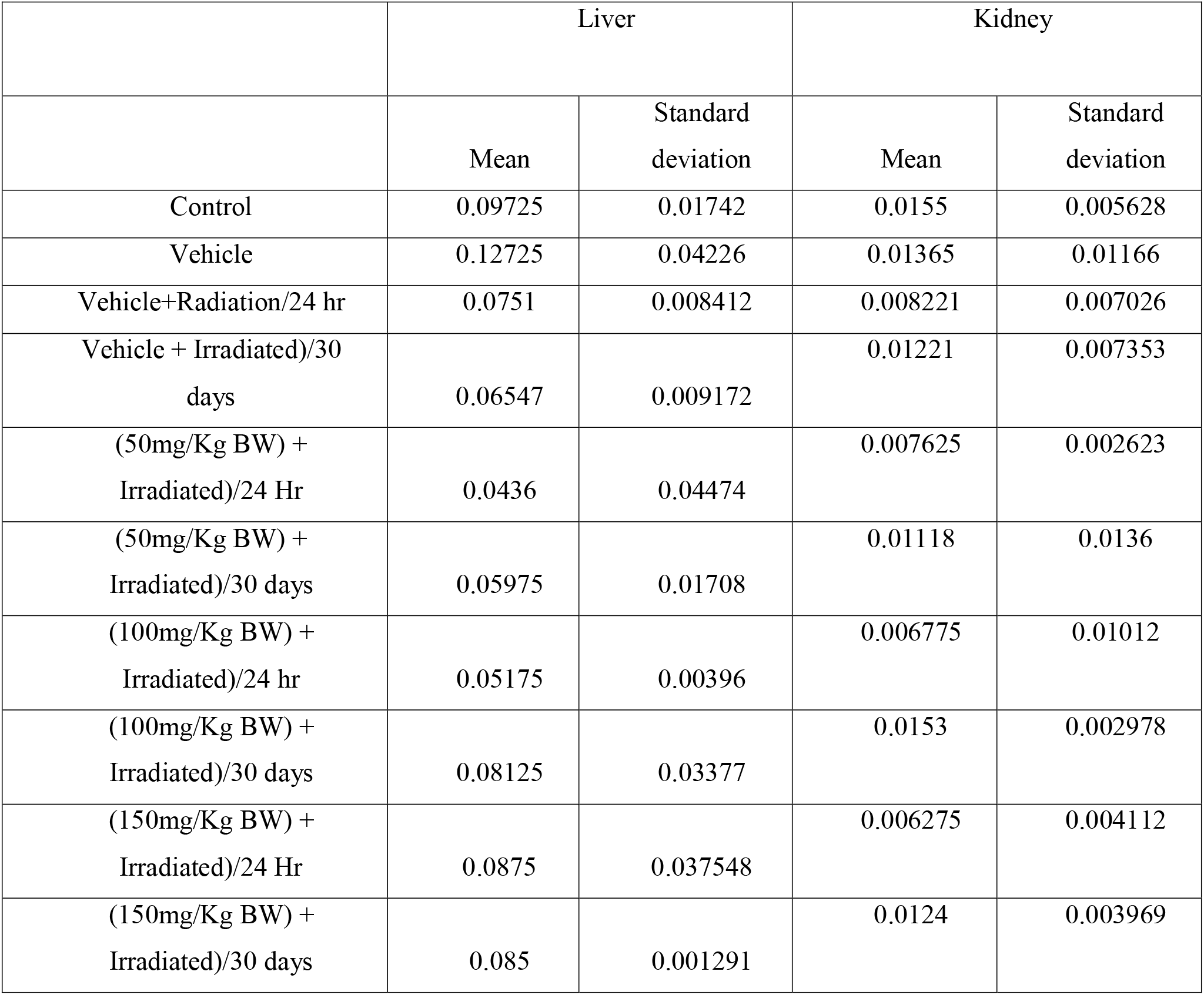
Modulatory effect of Omega 3 fatty acids on catalase activity (U/20) on radiation induced radiotoxicity in liver, kidney.

**Table 16:** Modulatory effect of Omega 3 fatty acids on NO scavenging activity (%) on radiation induced radiotoxicity in liver, kidney.

|  | Liver |  | Kidney |  |
| --- | --- | --- | --- | --- |
|  | Mean | Standard deviation | Mean | Standard deviation |
| Control | 16.185 | 3.156978 | 17.2 | 5.156 |
| Vehicle | 22.438 | 5.3442 | 18.02 | 7.464 |
| Vehicle+Radiation/24 hr | 11.2845 | 4.6857 | 13.18 | 8.992 |
| Vehicle + Irradiated)/30 days | 13.117 | 3.7381 | 15.25 | 9.585 |
| (50mg/Kg BW) +<br>Irradiated)/24 Hr | 13.3275 | 5.336461 | 13.51 | 1.464 |
| (50mg/Kg BW) +<br>Irradiated)/30 days | 14.0375 | 1.65527 | 18.34 | 1.187 |
| (100mg/Kg BW) +<br>Irradiated)/24 hr | 13.1675 | 3.718952 | 14.23 | 2.548 |
| (100mg/Kg BW) +<br>Irradiated)/30 days | 13.635 | 2.894641 | 17.4 | 2.151 |
| (150mg/Kg BW) +<br>Irradiated)/24 Hr | 13.03 | 2.866103 | 12.83 | 3.325 |
| (150mg/Kg BW) +<br>Irradiated)/30 days | 13.63 | 3.431229 | 18.62 | 1.171 |

**Table 17:** Modulatory effect of Omega 3 fatty acids on Total Antioxidant Capacity (ug/mg) in radiation induced radiotoxicity in liver, kidney.

|  | Liver |  | Kidney |  |
| --- | --- | --- | --- | --- |
|  | Mean | Standard Deviation | Mean | Standard deviation |
| Control | 64.3875 | 8.611217 | 44.08 | 12.24 |
| Vehicle | 61.8538 | 6.2868 | 45.21 | 14.16 |
| Vehicle+Radiation/24 hr | 49.32551 | 5.0847 | 39.21 | 13.74 |
| Vehicle + Irradiated)/30 days | 52.1274 | 6.00849 | 41.06 | 11.41 |
| (50mg/Kg BW) +<br>Irradiated)/24 Hr | 53.52 | 6.358587 | 38.58 | 1.674 |
| (50mg/Kg BW) +<br>Irradiated)/30 days | 58.4025 | 7.523956 | 35.48 | 5.814 |
| (100mg/Kg BW) +<br>Irradiated)/24 hr | 59.015 | 5.113813 | 31.91 | 3.605 |
| (100mg/Kg BW) +<br>Irradiated)/30 days | 57.775 | 8.342359 | 41.78 | 4.181 |
| (150mg/Kg BW) +<br>Irradiated)/24 Hr | 58.325 | 6.394956 | 39.79 | 4.604 |
| (150mg/Kg BW) +<br>Irradiated)/30 days | 57.3475 | 6.216173 | 42.41 | 4.899 |

### 3.5 Omega 3 fish oil enhance DNA repair in radiation induced toxicity in liver and kidney

DNA gel electrophoresis has been observed after radiation exposure with and without omega 3 fish oil. Liver has smear kind pattern even treated with omega 3 fish oil but normal band has been observed when DNA observed after recovery period of 30 days (Fig 4A, 4B and 4C). On the other hand, kidney shown less smear and more intact band even after exposure of electron beam radiation at post 24 hrs.

### 3.6 Omega 3 fish oil enhances the Wnt canonical pathway leads to inhibit the apoptotic event coupled with the DNA repair mechanism

Ku70 and Ku80 are two proteins responsible for the nonhomologous end-joining pathway (NHEJ), which joins DNA double-strand break. Also survivin is another protein which bind with caspase 3 and inhibit the apoptosis. In our study, Ku70 and survivin mRNA level in liver, kidney, increased when administered with omega 3 fish oil at 24 h post-radiation period (Figure 4(A), Table 3, Figures S3(A–D)). The mRNA level of Ku70 and survivin was more improved during the post-radiation period of 30 days compared to radiation alone when omega 3 fish oil was prior administered before radiation, indicating a poor recovery in repairing DNA and inhibiting apoptosis. In addition, damage to renal and hepatic irradiated cells was investigated with a DNA ladder (Figure 4(B)), with severe DNA damage at all doses over the post-24-h irradiation periods. Overall, agarose gel electrophoresis has shown that DNA shearing suggests apoptosis and necrosis.

## 4. Discussion

In our study we have found that Omega 3 fish oil has been a potent scavenger in neutralising the radiotoxicity at extent. Furthermore, the omega 3 fish oil enhances the enzymatic antioxidant level even at mRNA level. Radiation alone has elevated the level of Oxidative stress as well as decreased the enzymatic activity.

### 4.1 Wnt canonical pathway and omega 3 fish oil enhance cell survival during radiation

Previously, we explained how the Wnt canonical pathway in these events had shown its significant role in our study of post-radiation. High levels of ROS irrevocably damage cellular components, including its biomolecules, which further precede cell death to control the toxic threshold after IR is required to maintain ROS levels (Lento et al., 2014) It has been demonstrated that EPA and DHA limit pancreatic tumour cell development by interfering with the Wnt/-catenin signalling pathway and cause cell death by generating reactive oxygen species (ROS), activating caspase-8, or inducing autophagy(D’Eliseo & Velotti, 2016; Kyoung et al., 2011) Our study report shows that there is an upregulation in Wnt canonical pathway when administered with omega 3 fish oil suggesting protecting property of omega 3 fatty acids by underlying coupled and controlled molecular mechanism which executes radiation exposure via the Wnt canonical pathway by exploring the expression levels of its marker Lef1 and Axin2. The increase in mRNA level of Lef1, Axin2, and enzymatic antioxidants leads to activation of cell proliferation and cell survival, suggesting the effect of omega 3 fish oil in the Wnt canonical pathway during radiation. Our study suggests that omega 3 fatty acids induced Wnt activation can help protect the cells and control the undesirable apoptosis as our study observed recovery in the expression of its candidate molecules.

Additionally, the mRNA of enzymatic antioxidants has been improved in omega-3 fish oil-administered mice. Similar to this, (Hai et al., 2012)demonstrated that the Wnt canonical pathway protects stem/progenitor cells from radiation damage through the phosphorylation of the Wnt coreceptor LPR6, Wnt/b-Catenin signalling may also prevent the action of the enzyme glycogen synthase kinase 3b (GSK-3b) from occurring in the liver, kidney, and spleen(Cselenyi et al., 2008) Inhibitors of GSK-3b prevent radiation-induced apoptosis in the small intestinal epithelium and hippocampus neurons by blocking pro-survival transcription factors and promoting pro-apoptotic transcription factors (Thotala et al., 2010).

### 4.2 Omega 3 fish oil enhances Ku70 level suggesting improvemt in NHEJ pathway in mice

Ku70/Ku80 is the protein responsible for repairing DNA double-strand breakage in general via the NHEJ pathway (Davis & Chen, 2013). Our study suggests that the Omega 3 fish oil enhances Ku70 level suggesting improvement in NHEJ pathway resulting in protecting from DNA damage at extent. Recovery has been observed in the mice who were given omega-3 fish oil before the radiation. DNA lesions including abasic sites, oxidized bases, single-strand breaks and DSBs. Cells produce multiple ROS scavengers to defend themselves against such oxidative threats(Shiloh, 2014; Slupphaug, 2003). Other studies have also shown that high doses of radiation often lead to dominant lethal effects, point mutations, and chromosomal abnormalities (Arnon et al., 2001; Boudaı◻ffa et al., 2000) have shown that abundant low energy 1–20 eV secondary electrons (radical reaction products) play a crucial role in the nascent stages of DNA radiolysis. The ionization of nucleic acid bases has been identified as an initiating step of radiation damage and has been studied extensively with high energy resolution (Choi et al., 2008)

### 4.3 Omega 3 fish oil enhances DNA damage repair, coupled with the Wnt canonical pathway

Senescence, apoptosis and inflammation are caused by persistent DNA damage. According to a recent study by (Terada et al., 2002)EPA can prevent cell death induced by H2O2(Terada et al., 2002). Interestingly, the incidence of telomere shortening over 5 years in patients with coronary artery disease was shown to be inversely correlated with baseline blood levels of marine n-3 PUFAs in an intriguing prospective cohort study by (Farzaneh-Far, 2010). Telomeric attrition, a form of DNA damage, is a result of ROS that selectively target a specific sequence in telomeres. By preventing the telomere attrition brought on by ROS, Omega-3 PUFAs may slow down cell senescence. Additionally, Gray et al. showed using transgenic mice that poor DSB repair alters plaque phenotype and may be a factor in vulnerable plaques(Gray et al., 2015) Considering this, Omega-3 PUFAs may retard the development of atherosclerotic plaques and increase plaque stability in part by preventing DNA damage and the ensuing senescence of cells (Sakai et al., 2017) We previously explained two significant pathways involve DSB repair: homologous recombination (HR) and non-homologous end-joining (NHEJ). Ku70 inhibits cell proliferation, possibly by decreasing b-Catenin/Wnt signaling pathway during DNA damage (Budinger et al., 2002). We have observed that the Wnt canonical pathway’s candidate molecule and NHEJ candidate molecules are up-regulated at extent when mice are treated with omega 3 fish oil, suggesting omega 3 fish oil somehow trying to protect from cell death by enhancing the NHEJ and Wnt canonical pathway. The current study provides experimental evidence that omega-3 fish oil enhances the canonical Wnt-signaling pathway and mediates radiation-injured cells for repair.

### 4.4 Omega 3 fish oil suppresses Oxidative stress to an extent, a prominent supplement in reducing radiotoxicity and inducing/inhibiting combined Cascades of NHEJ, Wnt canonical, and intrinsic apoptotic pathways

Our study found an elevation in lipid peroxidation level and an increase in TAC level showing omega 3 fish oil supplementary factor in mice. Previous studies have shown a two-fold increase in LPO in the liver, kidney, brain, spleen, and testis (Kramer et al., 1991). TAC status shows the total antioxidant to fight out the ROS in cells. Hence, the increase in TAC shows the strengthening of the defence mechanism by omega-3 fish oil, which leads to cell protection. Superoxide dismutase and catalase are enzymatic antioxidants that break down H2O2 in the water; they are essential antioxidants in the defence mechanism (Bhatia & Manda, 2004). They work together as superoxide dismutase converts superoxide into peroxide, and catalase further converts peroxide into water. Higher catalase activity in irradiated mice indicates enzyme induction due to increased H2O2 formation (Weydert & Cullen, 2010)a natural adaptive response. The omega 3 fish oil administered mice showed a prominent recovery in oxidative stress at 24 h post-irradiation and 30 days post-irradiation, suggesting the omega 3 fish oil has antioxidant properties to some extent.

Similarly, a dose-dependent decrease in mRNA levels of SOD1 and Catalase has been observed, suggesting the recovery in enzymatic antioxidant activity. Nitric oxide scavenging activity reflects the defence mechanism by the cell to protect itself from nitric oxide free radicals. Nitric oxide free radicals are the byproducts further created by ROS. In this study, we observed the increased mRNA levels of enzymatic antioxidant gene by administration of Omega-3 PUFAs. Similarly, Sakai et al., 2017 showed HO-1 degrades heme and generates biliverdin, carbon oxide and iron. Biliverdin is converted to bilirubin by biliverdin reductase, and both of these bile pigments efficiently scavenge peroxyl radicals(Sakai et al., 2017). It has also been abbreviated as RNS, which stands for reactive nitrogen species. It is the first attempt to evaluate Nitric oxide scavenging activity in irradiated mice and improvement in nitric oxide scavenging activity by omega 3 fish oil. However, numerous studies have been done to evaluate Nitric oxide’s function in cell physiology (Jagetia & Baliga, 2004). NO is a crucial bioregulatory molecule with several physiological roles, including blood pressure control, neural signal transduction, platelet function, and antimicrobial and antitumor activities(Huang et al., 2007).

The present study found iNOS expression has been prominently improved dose-dependent when administered omega 3 fish oil prior to radiation, suggesting recovery of NOS activity leads to various pathophysiologies.

## Acknowledgments

The author wishes to thank Dr. Vishweshwara and HCG Bharath Cancer hospital staff for providing the irradiation facility. The author wishes sincere thanks to the Indian Council of Medical Research and Board of Research in Nuclear Sciences for providing us funding to carry out this work.

## Author contributions

Shashank Kumar conceived and designed the experiment, performed animal handling, harvesting of organs and germ cells, mRNA isolation, cDNA preparation, oxidative stress parameters, DNA ladder, Sperm morphology, viability, and motility. Shashank Kumar performed RT-qPCR analysis. Shashank Kumar performed the statistical analysis and prepared the manuscript. Suttur S. Malini corrected the manuscript. All authors read and approved the manuscript.

